# A Translational Preclinical Strategy for Chronic Spinal Cord Injury: Neuroprotective and Regenerative Potential of Botulinum Neurotoxin Type A combined with Muscle Atrophy Prevention via Electrostimulation

**DOI:** 10.64898/2026.03.23.713625

**Authors:** Valentina Mastrorilli, Siro Luvisetto, Veronica Ruggieri, Giada Raparelli, Luca Madaro, Lucia Amalia Paggi, Chiara Parisi, Francesca de Santa, Federica De Angelis, Annunziata D’Elia, Roberto Massari, Susanna Amadio, Ornella Rossetto, Valentina Vacca, Maurizia Caruso, Gianluca Sferrazza, Flaminia Pavone, Sara Marinelli

## Abstract

**Background:** Spinal cord injury (SCI) triggers persistent neuroinflammation, gliosis, neuronal loss, and demyelination, leading to motor deficits and neuropathic pain. Botulinum neurotoxin type A (BoNT/A) has shown anti-inflammatory and neuroprotective effects in acute SCI, but its potential in the chronic phase remains unclear. This study investigates whether combining BoNT/A with electrical muscle stimulation (EMS) enhances recovery in chronic SCI.

**Methods:** Adult mice with severe thoracic SCI (paraplegic) underwent EMS (30 min/day for 10 non-consecutive days starting 3 days post-injury) or no stimulation. Fifteen days after SCI, animals received a single intrathecal injection of BoNT/A (15 pg/5 μL) or saline. Functional recovery was assessed up to 60 days as well as in moderate and mild SCI mice, neuropathic pain onset and maintenance were evaluated. Spinal cord tissue was analysed for astrocytic and microglial morphology, neuronal and oligodendroglia survival, myelin protein expression, and in vitro effects on oligodendrocyte precursor cells (OPCs). The phenotype of hindlimb muscles was evaluated through morphological and gene expression analyses.

**Results:** EMS was able to counteract muscle atrophy and fibrosis, and when combined with BoNT/A, also denervation. Moreover, the combination restored hindlimb motor function in chronic SCI, whereas BoNT/A or EMS alone were ineffective. Neuropathic pain, a common comorbidity associated with SCI, was mitigated by BoNT/A treatment even when administered in the chronic phase. BoNT/A reduced astrocytic hypertrophy and excitatory synapse association and was associated with a morphology-based redistribution of microglial profiles toward a resting-like classification, decreased apoptosis, and increased neuronal and oligodendroglia survival. Myelin basic protein expression was significantly elevated in vivo. In vitro, BoNT/A promoted OPC differentiation into myelinating oligodendrocytes, increased process complexity, and upregulated Myelin basic protein, galactocerebroside C, proteolipid protein, and myelin oligodendrocyte glycoprotein under both proliferative and differentiating conditions. Cleaved SNAP25 colocalization with OPC confirmed direct BoNT/A internalization and activity.

**Conclusions:** BoNT/A exerts multi-cellular neuroprotective actions in chronic SCI, supporting neuronal and oligodendroglia survival, reducing neuroinflammation, enhancing remyelination and the combination with EMS promotes substantial recovery of muscle homeostasis within a permissive microenvironment shaped by early stimulation. Its efficacy depends on a permissive microenvironment achieved through EMS. These results provide strong rationale for the clinical evaluation of BoNT/A as a therapeutic strategy for chronic SCI.

## Background

Traumatic spinal cord injuries (SCI) represent a significant and growing public health challenge, with an estimated 0.9 million new cases reported annually and over 20 million people globally affected by SCI-related disabilities (1). These injuries commonly result from domestic accidents, occupational hazards, vehicular accidents, sports-related trauma, and interpersonal violence, leading to a spectrum of sensory and motor dysfunctions. Beyond these primary impairments, SCI is associated with various complications, such as neuropathic pain, gastrointestinal issues, spasticity, and reduced life expectancy, which collectively create a substantial economic burden, exceeding $4 billion annually in costs related to home care, adaptive housing, medical visits, and hospitalizations (United States estimation) (2, 3). Given this impact, there is an urgent need for therapies that can address not only the initial injury but also the long-term disability associated with SCI.

Regenerative medicine has made notable advances in treating SCI through neuroprotective agents, genetic interventions, and pro-regenerative biotechnologies (4). However, CNS injuries remain highly resistant to repair due to the complex structural and molecular characteristics of the CNS, particularly the inhibitory environment created by severe injuries like SCI. After SCI, vascular disruption, ischemia, and inflammation hinder axonal regrowth, while the formation of a glial scar serves as a physical and biochemical barrier to repair. This regenerative failure results from the lack of growth-promoting signals and the accumulation of inhibitory factors at the injury site, which together prevent neuronal recovery.

SCI progresses through acute, subacute, and chronic phases, with each stage presenting distinct therapeutic challenges. While early interventions may hold some potential during the acute phase, the chronic phase, beginning approximately four weeks post-injury, is marked by more profound and often irreversible losses in motor and sensory function. Chronic SCI is characterized by persistent neuroinflammation, glial scar formation, and muscle atrophy, all of which complicate any therapeutic approach. This inflammatory response, although useful for initial clearance of debris, can exacerbate tissue damage mediated by pro-inflammatory cytokines from activated microglia and infiltrating immune cells. Moreover, the degeneration of oligodendrocytes and subsequent demyelination further degrade the white matter, worsening functional outcomes (5,6).

In this context, botulinum neurotoxin type A (BoNT/A) stands out as a promising candidate for SCI treatment due to its unique biochemical and therapeutic properties. BoNT/A is a neurotoxin that specifically cleaves SNAP-25 (7), a crucial SNARE protein essential for neuroexocytosis (8). By blocking synaptic vesicle release, BoNT/A effectively modulates the secretion of neurotransmitters and inflammatory molecules, making it a valuable therapeutic agent across a spectrum of over 100 conditions, ranging from muscular to neurological disorders (9). Widely utilized in clinical SCI settings for its efficacy in treating spasticity, neurogenic bladder dysfunction, and neuropathic pain, BoNT/A has demonstrated a favourable safety profile with well-defined and manageable adverse effects, contributing to its reputation as a safe and effective treatment.

One of BoNT/A’s most beneficial attributes in clinical practice is its prolonged duration of action, often lasting several months due to its biochemical mechanism, which enables long-term effects from a single administration (10). This prolonged action supports sustained symptom relief, reducing the need for frequent treatments and improving patient compliance. Additionally, several studies (11) have uncovered broader pharmacological effects of BoNT/A, revealing its anti-inflammatory and neuroprotective properties that make it an appealing candidate for neuroinflammatory conditions. In preclinical SCI models, we have shown that BoNT/A administration during the acute phase (12,13) can promote motor recovery, decrease inflammation, and support axonal repair. These findings highlight its capacity to positively modulate the CNS environment, offering benefits beyond its established role in muscle tone and spasticity management. Given the high prevalence of SCI and the extensive demands of chronic care, we now focus on the therapeutic application of BoNT/A in the chronic phase of SCI, where intervention options remain extremely limited. For this purpose, we developed a murine model that mimics the chronic aspects of human SCI, including sustained neuroinflammation, muscle atrophy, and sensory-motor deficits (14,15). In this study, we implemented a muscle rehabilitation phase with electrostimulation to mitigate atrophy, followed by the administration of BoNT/A directly into the spinal cord. Our findings indicate that BoNT/A, even in the chronic phase, may reduce neuroinflammation, improve muscle tone, and promote remyelination, as evidenced by behavioural improvements and supportive histological and in vitro data.

The present study is part of a repurposing pharmacological R&D program that led to the identification of new mechanisms of action of BoNT/A and a new route of administration of the drug, promoting its future use as innovative experimental medicine for the treatment of SCI patients. To this end, a translational preclinical strategy, in line with regulatory guidelines, has been conducted, and considering the promising results obtained, we are currently seeking approval for a clinical trial investigating the BoNT/A’s potential in this therapeutic field (16).

Our findings not only underscore BoNT/A’s transformative role in neuroinflammation and remyelination but also paves the way for translating these preclinical findings into clinical practice. Our aim is to contribute a novel, multidisciplinary approach to SCI management that could improve quality of life and clinical outcomes for individuals living with chronic SCI.

## Methods

### Animals

Four to six-month-old CD1 female mice (EMMA Infrafrontier, Monterotondo - Italy) were used. Weighing about 35–40 g at the beginning of the experiments, mice were housed in groups of 4 or 8 in standard cages under a 12/12 h light/dark cycle (7:00 a.m.–7:00 p.m.) with food and water available ad libitum. Before being used for any experimental setup, the animals were acclimated in our animal facility for at least 30 days and, similarly, thirty minutes before surgery they were placed in the experimental room for the acclimatization. Testing was done by blind investigators as for treatment groups (as better described in “Randomization, allocation concealment and blinding” section). Care and handling of mice were in accordance with the guidelines of the Committee for Research and Ethical Issues of IASP (PAIN® 1983, 16, 109–110). Experimental protocol for in vivo procedures was approved by the Italian Ministry of Health (protocol code 122/2019PR) The number of animals utilized for each experiment and group is reported in the figure legend.

### Surgery

To induce severe SCI, animals were deeply anesthetized using either a 1:1 mixture of Rompun (Bayer, 20 mg/ml; 0.5 ml/kg) and Zoletil (100 mg/ml; 0.5 ml/kg), or connected to an inhalation anaesthesia system delivering isoflurane (1–3%) in a continuous oxygen flow (2 L/min) through a nose cone. After shaving the back with an electric razor, the skin was disinfected with Betadine and incised to expose the vertebral column (12,13). Animals were then secured in a stereotaxic frame equipped with spinal adaptors and connected to the PinPoint Precision Impactor Device (Stoelting; Hatteras Instruments Inc., Cary, NC, USA). Core body temperature was maintained at 37 °C throughout the procedure.

Spinal cord contusion was induced without laminectomy at thoracic levels T9–T11 (Figure S1), using the following parameters: impactor tip #4 (middle, round, flat), velocity 3 m/s, depth 5 mm, and dwell time 800 ms. Following surgery, animals were monitored daily for postoperative complications. Postoperative care included housing under red light for the first 24 hours, subcutaneous injection of betamethasone (1 mg/kg), and provision of water-based gel and softened food in the cage to support hydration and nutritional intake. Bladders were manually expressed twice daily until spontaneous voiding was restored.

Potential outliers were identified through analysis of impact parameters recorded by the PinPoint software. Behavioral and observational criteria were also used to assess variability in injury severity across groups. It is important to note that not all animals subjected to “severe” injury parameters developed complete hindlimb paralysis, likely due to individual differences in spinal cord compression caused by the underlying vertebral structure. Although most experiments were conducted on animals with severe injuries (Basso Mouse Scale (17)- BMS score 0–3), a subset of animals with moderate (BMS 4–6) or mild (BMS 7–9) injuries was also included in specific behavioral assays to evaluate the potential effects of treatment on neuropathic pain.

### Electrical Muscle Stimulation

In preliminary experiments, we also tested a passive locomotor rehabilitation protocol using treadmill training (see Supplementary Methods and Figure S2 for details). This approach was initially chosen with the rationale of exploiting residual spinal locomotor circuits (central pattern generators) (18) that might be recruited by repetitive stepping-like movements. However, despite the progressive increase in training speed and duration, treadmill training failed to promote motor recovery in severely injured animals, as BMS scores remained unchanged over the 60-day observation period. Based on this lack of efficacy, we focused on EMS as a more effective strategy to prevent muscle atrophy and to create a permissive environment for subsequent BoNT/A treatment

Seventy-two hours after surgery, animals were randomly assigned to stimulated or non-stimulated groups. Mice in the stimulated group underwent hind paw EMS for 30 minutes per session, across 10 non-consecutive days (EMS protocol Figure 1B). Stimulation parameters were: pulse duration 160 μs, frequency 60 Hz, delivered using a Tesmed® stimulator (Shenzhen Roundwhale Technology Co., Ltd), similarly to protocols utilized in murine models (19). Smallest (4x4 cm) transcutaneous electrodes patches were utilized and hindlimbs entirely enveloped (Video S1). To optimize electrical conductivity, fur was shaved from the hind paws and an electroconductive gel was applied prior to each session. Stimulation was performed in a dedicated experimental cage where animals were temporarily housed and allowed to move freely; no physical restraint was required. Only mice with severe SCI were included in the EMS group.

**Figure 1.**
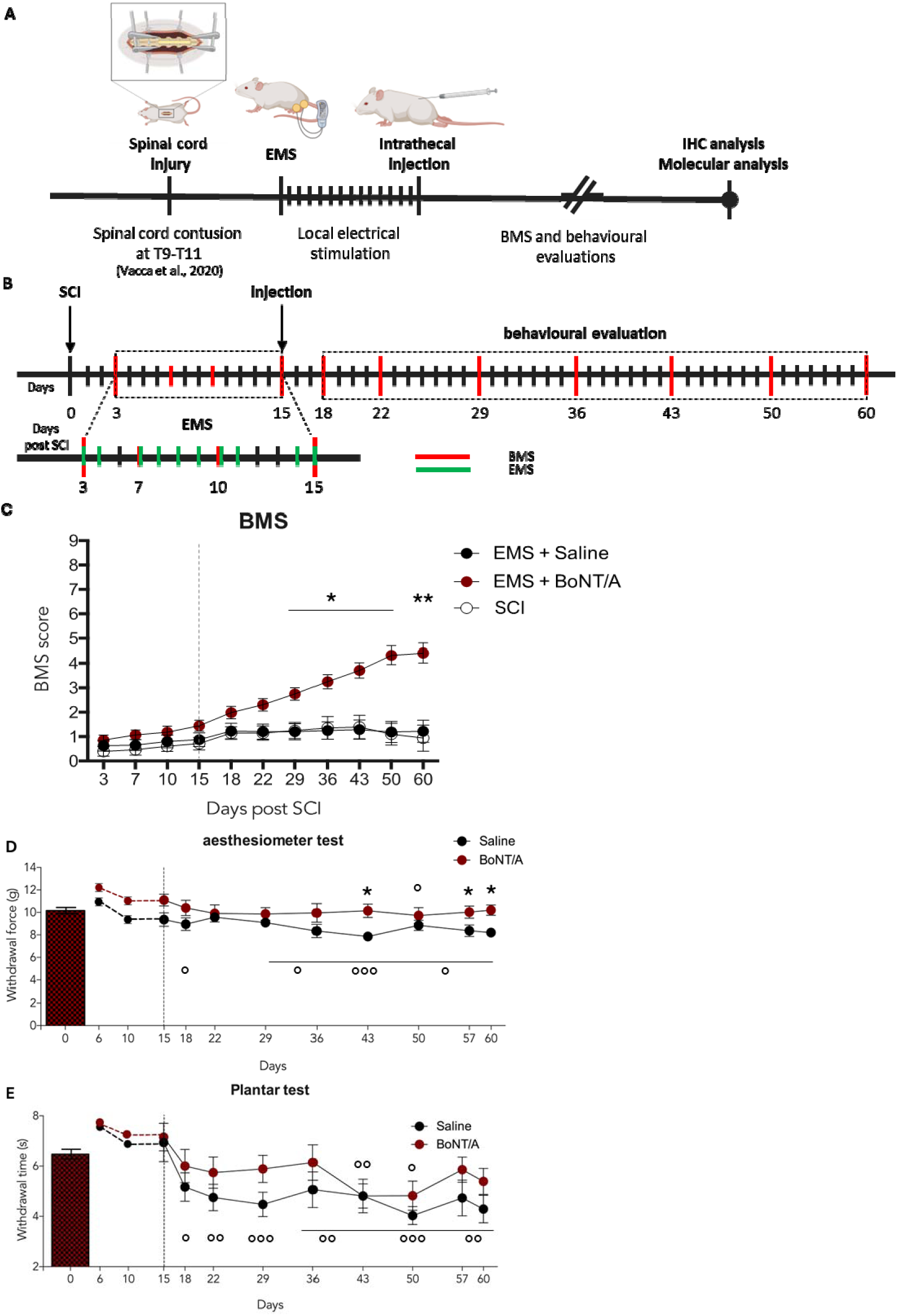
Effects of combination of EMS and BoNT/A on motor recovery. **A**) Schematic representation of the experimental paradigm combining a rehabilitative protocol with spinal BoNT/A injection. The study design integrates an EMS protocol, aimed at preventing muscle atrophy, with a single intrathecal injection of BoNT/A administered 15 days after spinal cord injury. **B)** Detailed timeline of the EMS protocol, with green line indicating the scheduled days of EMS. Red lines mark the days on which behavioral assessments were performed**. C)** Basso Mouse Scale (BMS) evaluation of motor function in mice with severe spinal cord injury (BMS score 0–3). Groups included: untreated SCI animals (SCI), animals receiving EMS combined with intrathecal saline injection 15 days post-injury (EMS+Saline), and animals receiving EMS combined with BoNT/A (EMS+BoNT/A). Dotted line indicates the day of treatment. No differences before treatment were revealed between groups. After treatment (from day 18 – D18) repeated measures ANOVA revealed a significant effect of treatment (F_2,29_ = 6,137 p = 0.0006), time (F_6,12_ = 4,529, p = 0.0003), and treatment × time interaction (F_12,174_ = 4,431, p = <0.0001).Tukey-Kramer post hoc analysis showed significant differences at days 29, 36, 43, 50, and 60 (*=p <0.05; **=p<0.005 ) between EMS+BoNT/A and both EMS+ Saline and NO EMS + Saline groups (N =10 EMS+Saline group; N=16 EMS + BoNT/A; N=6 No EMS + Saline). **D)** Mechanical allodynia was assessed in mildly to moderately contused mice using the Dynamic Plantar Aesthesiometer (force expressed in grams). A repeated measures ANOVA conducted between days 18 and 60 post-SCI revealed a significant main effect of treatment (F_1,34_ = 8.278, p = 0.0069), with no significant effects for time or for the treatment × time interaction. Each group included 9 animals (N = 18 values per time point, considering both hind paws were analyzed separately). No significant differences were detected between groups or for the group × time interaction during the pre-treatment phase (repeated measures ANOVA). Post hoc Tukey-Kramer analysis revealed significant differences between BoNT/A- and saline-treated mice at D43, D57, and D60 (*p < 0.05). Paired t-tests versus baseline (D0, pre-SCI) (°=p<0.05; °°=p<0.005; °°°=p<0.0001) showed a significant reduction in mechanical threshold in BoNT/A-treated mice at D50 (tLL = 2.122, p = 0.048), whereas saline-treated mice displayed a significant development of mechanical allodynia at D18 (tLL = 2.11, p = 0.050), D29 (tLL = 2.297, p = 0.034), D36 (tLL = 3.017, p = 0.0078), D43 (tLL = 5.773, p < 0.0001), D50 (tLL = 2.198, p = 0.0421), D57 (tLL = 4.041, p = 0.0008), and D60 (tLL = 2.97, p = 0.0086). **E)** Thermal hyperalgesia was evaluated in mildly to moderately contused mice using the Plantar Test. During the pre-treatment phase, no significant differences were observed between groups or with respect to baseline values. Repeated measures ANOVA conducted for the post-treatment period (D18–D60) did not reveal significant effects for treatment, time, or treatment × time interaction, although treatment approached statistical significance (F_1,36_ = 3.539, p = 0.068). Paired t-tests versus baseline (D0, pre-SCI) (°=p<0.05; °°=p<0.005; °°°=p<0.0001) showed a significant decrease in thermal withdrawal latency (indicative of thermal hyperalgesia) in BoNT/A-treated mice at D43 (tLL = 3.175, p = 0.005) and D50 (tLL = 2.182, p = 0.0419). In saline-treated animals, significant thermal hyperalgesia compared to baseline was observed at D18 (tLL = 3.929, p = 0.0011), D22 (tLL = 4.682, p = 0.0002), D29 (tLL = 5.420, p < 0.0001), D36 (tLL = 2.906, p = 0.0035), D43 (tLL = 4.577, p = 0.0003), D50 (tLL = 6.730, p < 0.0001), D57 (tLL = 4.750, p = 0.0002), and D60 (tLL = 4.720, p = 0.0002). Each time point included 20 measurements for BoNT/A (10 animals × 2 hind paws) and 18 for saline (9 animals × 2 hind paws).

A small number of severely injured animals (n = ) were assigned to the non-stimulated group. This decision was made in accordance with the 3Rs principle (Replacement, Reduction, Refinement), as the SCI protocol induces a high degree of suffering and is associated with prolonged impairment over the 60-day experimental period. Based on our previously published data (12, 13, 14), animals subjected to severe SCI without EMS show no spontaneous functional recovery and in fact exhibit progressive deterioration in general health and behavior. Therefore, only a minimal number of animals were allocated to untreated conditions to reduce unnecessary distress while preserving scientific validity.

### Drug administration

Fifteen days after SCI, animals were re-anesthetized using inhalation anaesthesia as previously described. A single intrathecal injection of either BoNT/A (15 pg in 5 μL; 150 kDa purified neurotoxin gently gifted by Prof. C. Montecucco, University of Padua) or saline (0.9%, 5 μL) was administered using a 10 μL Hamilton® syringe (30-gauge needle; Biosigma, Cona [VE], Italy).

As previously demonstrated, BoNT/A is capable of retrograde axonal transport (11,20–22). To minimize the risk of systemic diffusion potentially associated with ischemic events at the site of contusion, the injection was performed caudally, between the lumbar vertebrae L4 and L5.

### Randomization, allocation concealment and blinding

All mice assigned to EMS underwent rehabilitation and behavioral testing by Experimenter #1. At the end of the EMS phase, Experimenter #2 prepared either BoNT/A or vehicle solutions and labelled each vial with an alphanumeric code (distinct code per vial); the content was concealed to all other operators. Subsequently, Experimenter #3, who had not participated in the rehabilitation/testing phase, randomly selected and injected the solution from vial ‘1’ or ‘2’ under blinded conditions. Following injections, animals, now randomly allocated to treatment, were re-tested and harvested by Experimenter #1, still blinded to group identity. Harvested tissues were then distributed to collaborators for the various assays (histology, imaging, WB, qPCR), each sample identified only by its code; codes were revealed after data lock and statistical analysis. This three-operator workflow ensured allocation concealment, blinding of outcome assessors, and randomization of treatment assignment.

### High-Resolution In Vivo Micro-CT Imaging for Quantitative Analysis of Spinal Skeletal Structures

Micro-computed tomography (micro-CT, Figure S1) was employed to obtain high-resolution in vivo datasets of spinal skeletal morphology using a MILabs Hybrid OI/CT imaging system (MILabs, Houten, The Netherlands). Mice were anesthetized via inhalation of isoflurane (1–3%) in a continuous oxygen flow (2 L/min) delivered through a nose cone, ensuring stable immobilization during image acquisition. Scans were performed in accurate acquisition mode, comprising 720 angular projections over a 360° rotation arc. The X-ray tube was set to 35 kV (0.43 mA), with an exposure time of 200 ms per projection. All acquisition parameters were held constant across experimental subjects to ensure reproducibility and facilitate comparative analysis. No respiratory or cardiac gating was applied. Each imaging session lasted approximately 5 minutes, generating volumetric datasets with high isotropic resolution suitable for detailed structural evaluation of vertebral components.

### Three-Dimensional Reconstruction and Skeletal Segmentation

Projection data were reconstructed using the proprietary MILabs software, producing volumetric datasets with isotropic voxel size of 80 μm. The reconstructed images were exported in DICOM format and analysed using Imalytics Preclinical image analysis software (Gremse-IT GmbH, Germany). Grayscale values were calibrated to Hounsfield units to allow standardized interpretation. Three-dimensional segmentation and rendering of the spinal cord were subsequently performed to isolate and quantify vertebral and adjacent skeletal structures within the thoracolumbar region.

### Behavioural tests

Basso Mouse Scale (BMS). For injured animals, the hindlimb locomotor function was assessed in an open field. The BMS score ranges from 0 to 9, where 0 indicates complete paralysis and 9 normal movement of the hindlimbs (17). Performances of the left and right hindlimbs were averaged to obtain the BMS score. Mice were tested for hindlimb functional deficits at D3, D6, D10, D15 (injection timepoint), D18, D22, D29, D36, D43, D50, D57 and D60 after SCI. Severe, moderate and mild injured mice were included in this test. Mild and moderate contused animals were assigned to neuropathic pain evaluation tests.

Based on preliminary experiments showing that BoNT/A administered alone in the chronic phase did not restore motor function in this severe model (Supplementary Figure 4), subsequent mechanistic analyses were focused on EMS-conditioned cohorts to evaluate therapeutic efficacy within a functionally permissive context.

### Mechanical and thermal Allodynia: Dynamic Plantar Aesthesiometer and plantar test

Mechanical allodynia was tested using a Dynamic Plantar Aesthesiometer (Model 37,400, Ugo Basile Srl, Comerio, Italy) as previously described (23).

For habituation, mice were placed 30′ before test in the experimental room and in testing plastic cages with a wire net floor 5-10 min before the experiment. Each testing day, the withdrawal threshold of hind paws was taken as the mean of 3 consecutive measurements per paw.

Thermal hyperalgesia was tested using an automatic plantar test instrument (Plantar Test, Ugo Basile, Comerio, Italy). For the thermal hyperalgesia test, a cut-off time of 15 s was imposed to avoid damage to the hind paw skin tissue.

Mechanical and thermal threshold were measured at D3, D6, D10, D15 (injection timepoint), D18, D22, D29, D36, D43, D50, D57 and D60 after SCI and tests were conducted 1 hour spaced-apart. For each mouse, two values of mechanical and thermal threshold were obtained because the two hind paws, the right and the left, can develop different degrees of neuropathic pain. At each testing day, threshold values were averaged from 3 consecutive measurements per hind paw.

### RNA analysis by quantitative PCR

Total RNA was extracted from frozen gastrocnemius (GA) muscle tissue using Tri-reagent (Zymo Research) following the manufacturer’s instructions. RNA concentration and purity were assessed with a NanoDrop ONEc spectrophotometer (Thermo Scientific). First-strand cDNA was synthesized from the extracted RNA using the PrimeScript RT Reagent Kit with gDNA Eraser (Takara) according to the manufacturer’s protocol. Quantitative real-time PCR was performed on a QuantStudio 7 Flex system (Applied Biosystems, Thermo Scientific) using the ExcelTaq 2X Fast Q-PCR Master Mix (SYBR, ROX) (Smobio).

Gene-specific primer sets used for real time PCR were:

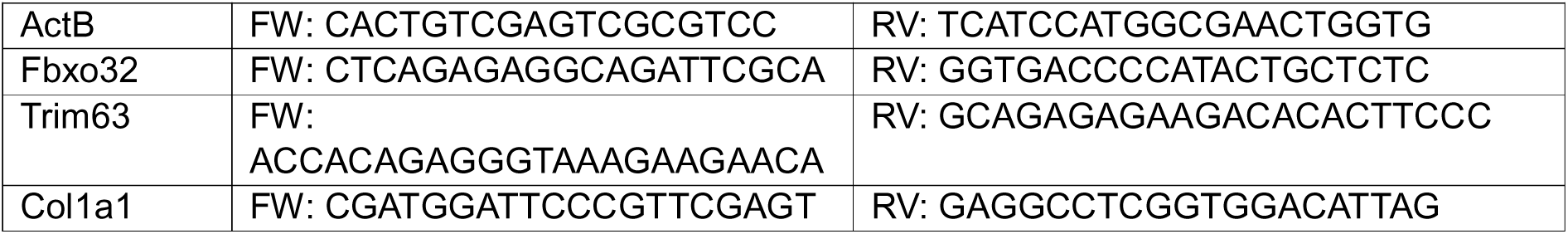

### Histological staining

#### Muscle Immunofluorescence staining

GA muscle cryosections (10 μm) were fixed in 4% paraformaldehyde (PFA; MilliporeSigma, P6148) for 10 minutes or permeabilized with 100% acetone for 1 minute at room temperature. Sections were blocked for 1 hour in PBS containing 4% bovine serum albumin (BSA; MilliporeSigma, A7030-100G). Primary antibodies were diluted in block solution and applied overnight at 4°C: anti-synaptophysin, anti-laminin and anti-caveolin-3 as specified in Table 1. After washing with PBS, sections were incubated with species-appropriate secondary (Table 1). Acetylcholine receptors (AChRs) were visualized using fluorescently labelled bungarotoxin (Table 1). Nuclear staining was performed with DAPI in PBS for 5 minutes. Finally, sections were washed in PBS and mounted in glycerol (3:1 in PBS). Each muscle was separately analysed. Images were acquired using a Zeiss confocal microscope and processed with ImageJ software.

**Table 1.**
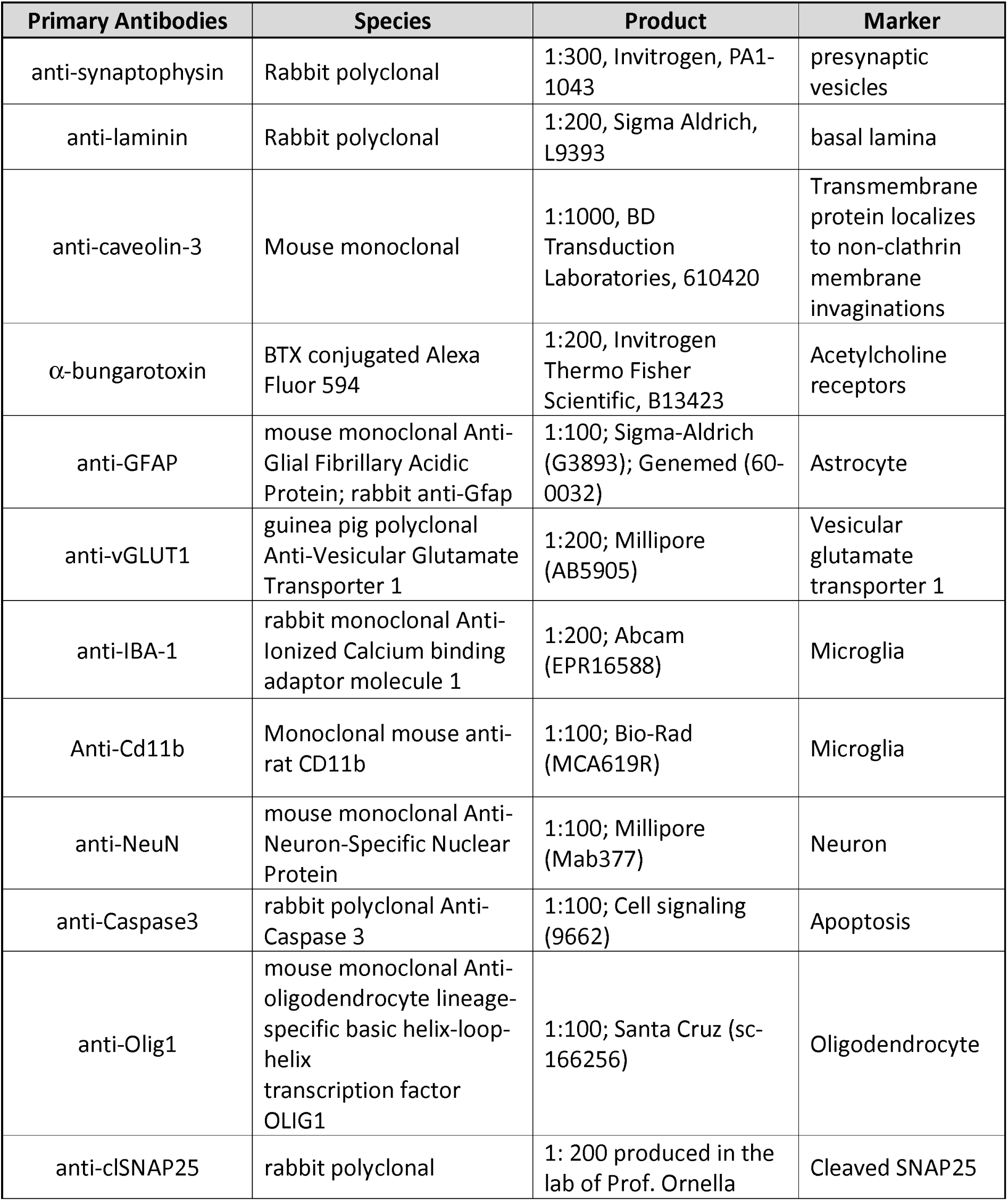
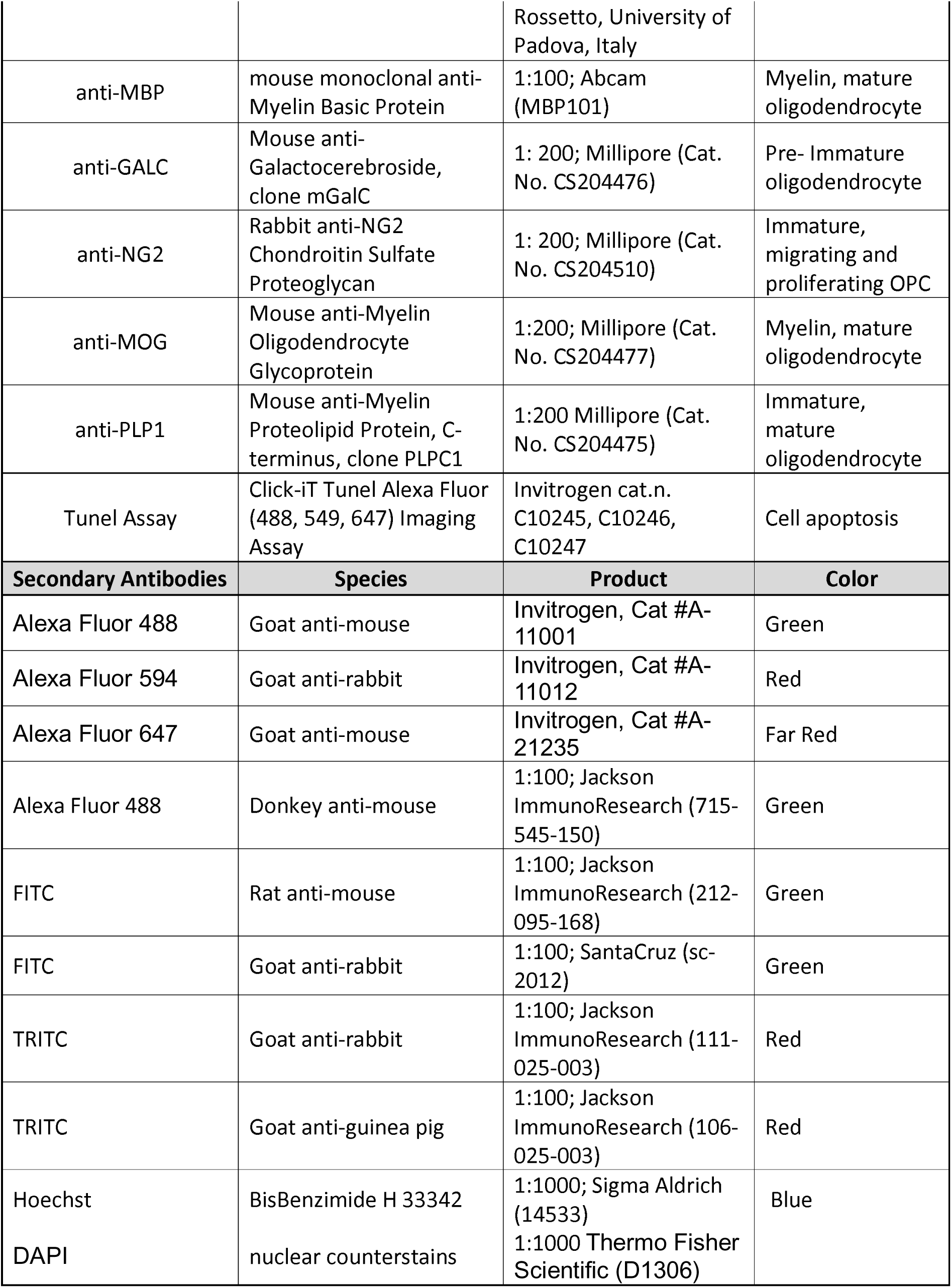
List of primary and secondary antibodies.

#### Sirius Red staining

GA muscle cryosections were fixed for 1 hour at 56°C in Bouin’s solution (Sigma-Aldrich, Cat #HT10132), then stained for 1 hour in Picro-Sirius Red solution (Direct Red 80 Cat #365548, Sigma-Aldrich) protected from light. Sections were briefly washed in acidified water (0.5% vol/vol), dehydrated in 100% ethanol, cleared in 100% toluene, and mounted with EUKITT mounting medium (Sigma-Aldrich, Cat #03989). Images were acquired with a Zeiss Imager.A2 microscope.

#### Haematoxylin and eosin (H&E) staining

**GA** sections were fixed in 4% PFA for 10 minutes, washed in PBS, then stained with haematoxylin (Sigma-Aldrich, Cat #HHS32) for 12 minutes and eosin (Sigma-Aldrich, Cat #HT110332) for 30 seconds. Sections were dehydrated in ethanol and mounted with EUKITT mounting medium (Sigma-Aldrich, Cat #03989).

#### Immunohistochemistry of spinal cord tissues

60 days post-SCI, three or four mice from each experimental group (except for the non-stimulated, untreated SCI group, for which only two/three animals were available) were sacrificed for immunohistochemistry and perfused with saline 0.9% followed by 4% paraformaldehyde in phosphate buffer saline (pH 7.4). Spinal cords were then collected and kept for 48 hours in PFA atL4 °C, then cryo-protected overnight in sucrose dissolved at 30% in PBS 1X and finally cryopreserved atL−L80 °C. Slicing of the spinal cord was carried out by embedding the tissues in Tissue-Tek OCT (Sakura, Torrence, CA, USA) and sections of 40μm thickness were collected. For double immunofluorescence staining, different sections were incubated for 48 hours at room temperature with primary antibodies (Table 1) in Triton 0.3%. Sections were then washed in PBS and incubated for 2 hours, at room temperature, with secondary antibodies (Table 1). Sections were again washed in PBS and incubated for 10 min with Bisbenzimide (Hoechst 33258, 1:1000, Jackson ImmunoResearch, Cambridge House, Ely, UK) to stain nuclei. Sections were finally mounted on glass slides with glycerol 3:1 in PBS.

#### Confocal microscopy and quantification of immunoresponsivity

Immunostained sagittal spinal cord sections were imaged using a TCS SP5 laser scanning confocal microscope (Leica Microsystems). All acquisitions were performed in sequential scanning mode to eliminate cross-channel bleed-through. Both low (5x and 20x objectives) and high (40x and 63x objective) magnification images were acquired and processed using I.A.S. software (Delta Systems, Italy).

To compare cell counts obtained from 63x and 40x objectives on an equivalent field basis, 63x counts were scaled to match the larger tissue area captured at 40×. Because the 40x field of view is 2.476-fold larger than the 63× field (150,156 µm² vs 60,657 µm²), 63x counts were corrected using:

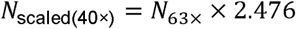

This area-based upscaling allows direct comparison between datasets acquired at different magnifications.

Quantitative analyses were carried out with Fiji/ImageJ software (National Institutes of Health, USA). NeuN-, and TUNEL-positive nuclei were automatically counted. TUNEL-positive nuclei were quantified using ImageJ. After background subtraction and signal saturation to enhance nuclear circularity, images were converted to binary masks. Cell detection was then performed with the “Analyze Particles” tool, applying a minimum size threshold of 50 pixels to prevent residual background artifacts or very small nonspecific elements from being counted as cells (n per each experiment/group and number of slices utilized are reported in figure legend), and group means were subsequently calculated.

Astrocytic and microglia distribution and morphology were analysed on confocal images of GFAP or IBA1/Cd11b, respectively, immunofluorescence acquired from spinal cord sections encompassing the lesion epicenter (T9–T11), perilesional regions (T7–T8 and T12–T13), and the epicenter-scar area. Each image was converted to 8-bit grayscale and binarized using the Otsu thresholding algorithm to generate a binary mask.

The total GFAP- or Iba1/Cd11b positive area was measured using the “Analyze Particles” function, expressing the value in pixels. The *size* and *circularity* parameters were set to discriminate cell bodies from processes divided from the soma. Specifically, a range of 200–∞ pixels (circularity 0–1) was used to quantify the total area occupied by GFAP-positive elements, including detached processes, while a more restrictive range of 400–∞ pixels was applied to estimate the number of astrocytic cell bodies. For microglia, since its small size, a range of 50–∞ pixels (circularity 0–1) was used to quantify the total area occupied by Iba1/Cd11b-positive elements, including detached processes, while a more restrictive range of 100–∞ pixels was applied to estimate the number of microglia cell bodies.

In the scar region, characterized by a dense and irregular GFAP network or cystic cavities devoid of cell bodies, only the total GFAP-positive area was measured. Representative images showing the original acquisition, binary mask, and segmented overlays are reported in the Supplementary Figure S3.

Fluorescence intensity for Caspase-3 and Olig1 was quantified using the RGB method, which converts red, green, and blue pixel values to brightness levels. In addition, signal quantification and colocalization within defined regions of interest were assessed using integrated density measurements and Manders’ colocalization coefficient.

#### Morphometric and Sholl analysis

Morphometric analysis was performed on high-resolution images acquired using the following confocal parameters: 40x or 63x objective, 3x digital zoom, 1024L×L1024 frame size, and 10 Hz scanning speed. Each image was converted to binary format to generate digital silhouettes of astrocytes or microglial cells, allowing for the identification and measurement of cellular outlines. Activated microglia were quantified using a previously established classification method (12).

Briefly, based on morphological features such as circularity and process length, microglia were categorized into distinct activation states: ramified (resting microglia), hyper-ramified (bushy or hyperactive microglia), and unramified/amoeboid (fully activated microglia). Cell classification was guided by representative reference images for each phenotype.

Sholl analysis was conducted on IBA1⁺/Cd11b^+^ microglia, GFAP⁺ astrocytes, and differentiated oligodendrocyte precursor cells (OPCs) to assess the complexity of process arborization (quantified as the number of intersections between microglial processes and concentric circles at increasing radial distances from the soma). Cell diameter was defined as the maximum radial distance from the soma to the tip of the longest projection. Concentric circles were drawn at 5 μm intervals from the cell soma, and the number of intersections per radius was quantified. This analysis enabled the evaluation of branching complexity as a function of radial distance from the cell body, following the method described by Paes-Colli et al. (24). Quantification was performed using the Sholl Analysis plugin in Fiji/ImageJ (NIH, Bethesda, MD; developed by Wayne Rasband), according to the developer’s instructions.

These data were used to evaluate microglial, astrocytic, and OPC process complexity and branching patterns in response to spinal cord injury.

#### Western Blot Analysis

Total proteins were extracted from spinal cord in RIPA buffer ((50 mM Tris-HCl pH 8.0; 150 mM NaCl; 1mM EDTA pH 8.0; 1% Triton; 0,1% SDS; 1% NaDOC) supplemented with protease and phosphatase inhibitors (5 mg/ml Aprotinin; 5 mg/ml Leupeptin; 5 mg/ml Pepstatin; 1 mM PMSF; 10mM NaF; 200 mM Na_3_VO_4_; 500 mM β-Glycerophosphate) by manual mincing of tissues with a plastic pestle, followed by sonication (microtip, power 2, 10 + 10 seconds). Then the samples were rocked on a wheel (20-30 minutes, +4°C) and centrifuged (12000 rpm, 15 minutes, +4°C), to remove the tissue debris and stored at -80°C. The protein concentration was detected by measuring 595 nm absorbance after staining with Bradford dye (BioRad 5000006). 10-25 μg of proteins was run on polyacrylamide gels and transferred electrophoretically to 0.45 μm nitrocellulose membranes (BioRad 162-0115) using the wet system from Bio-Rad. After blocking with 5% non-fat dry milk for 1 h, membranes were incubated at +4°C overnight with the primary antibodies: - NMDAε2 receptor (1:1000, ELK Biotechnology ES5659), - GABA-A receptor α2 (1:1000, ELK Biotechnology EA285), - MBP (1μg/ml; Abcam ab62631), GAPDH (1:5000, Invitrogen MA5-15738) and Actin β (1:5000, Invitrogen MA1-744); anti-GFAP (1ug/ml, Invitrogen #14989282 Mouse Monoclonal); anti GLUT1: SYSY (1:1000, Synaptic Systems #135 302 Rabbit Polyclonal ); anti EAAT1 (GLAST-1)(1:5000, Millipore #AB1782 Guinea Pig Polyclonal ).

After 3 washes with PBS 0,01% Tween, membranes were incubated with secondary antibodies (donkey anti-mouse IgG-HRP 1:10000, Jackson ImmunoResearch 715-035-150; donkey anti-rabbit IgG-HRP 1:10000, Jackson ImmunoResearch 711-035-152; anti Guinea Pig: Millipore #AP193P 1:5000) for 1h at room temperature and then washed with PBS 0,01% Tween.

Immunoreactivity was determined using the enhanced chemiluminescence luminol reaction and revealed by Chemi-Doc Imaging System (BioRad). Densitometric analysis was performed using Image J software.

#### In vitro oligodendrocyte cell cultures and BoNT/A treatment

Rat Glial Precursor Cells (GPCs), also referred to as Oligodendrocyte Progenitor Cells (OPCs) (Gibco, N7746100) were thawed and seeded at a concentration ranging from 1x10^5 to 2.5x10^5 cells/cm2 in complete OPC growth medium (KnockOut™ D-MEM™/F-12, 2% StemPro™ NSC SFM™ Supplement, 20ng/mL EGF, 20ng/mL bFGF, 2 mM GlutaMAX™-I and 10 ng/mL PDGF-AA) on poly-L-ornithine (20ug/mL) coated tissue culture plates, or coverslips.

Complete OPC medium was changed every two days to maintain undifferentiated proliferating cells. Otherwise, to induce spontaneous oligodendrocyte differentiation and maturation, OPC medium was replaced with oligodendrocyte differentiation medium (KnockOut™ D-MEM™/F-12, 2% StemPro™ NSC SFM™ Supplement, 2% FBS, 2 mM GlutaMAX™-I) and replaced with fresh medium every day.

To characterize the responsiveness of OPCs to BoNT/A, OPCs and differentiating oligodendrocytes were treated with vehicle or BoNT/A (10pM) for 48 h.

To assess cell viability cells were detached using StemPro™ Accutase™ Cell Dissociation Reagent (Gibco), stained with trypan blue and counted using a hemocytometer.

#### Immunofluorescence staining of cells

OPCs and differentiating oligodendrocytes were washed with phosphate buffered saline (PBS) and fixed using 4% PFA for 15 minutes, followed by permeabilization with PBSL+L0.1% TRITON X-100, blocked in blocking buffer (3% bovine serum albumin—BSA—in PBS), and subjected to incubation overnight at 4L with primary antibody. Then cells were washed 3X for 5 minutes with PBS and incubated with fluorescence-labeled secondary antibody in the dark at 37L for 30–45 minutes. After washing 3X with PBS, cells were incubated with DAPI solution (3 ng/mL) for 5 minutes, rinsed with PBS and coverslips mounted with fluoromount (Sigma Aldrich) on microscope slides. The list of primary and secondary antibodies is reported in Table 1.

#### Statistical analysis

The groups’ size for in vivo experiments was calculated by implementing a power analysis (Gpower 3.1). For locomotor recovery (BMS score; 11 repeated measures across 3 groups), sample size was calculated assuming effect size f = 0.25, α = 0.05, 1–β = 0.95 and indicated that 27 animals were required. For neuropathic pain tests (aesthesiometer and plantar test), repeated-measures ANOVA was performed separately for pre-treatment (4 measures including baseline) and post-treatment (8 measures). Power analyses with effect size f = 0.25, α = 0.05, 1–β = 0.80 yielded required sample sizes of 24 (pre-treatment) and 16 (post-treatment). The number of mice used is reported in the figure legends. With the regard of immune- and biochemical experiments, sample size was estimated according to previous experience, using the models described. Experimental data were expressed as mean ± SEM. Group comparisons were conducted by one-way or two-way ANOVA for repeated measures or by Student’s t-test. Post hoc comparisons were made with Tukey-Kramer tests (statistical significance at p < 0.05). Data analysis was performed by StatView SAS (version 5.0, Cary, NC, USA) or Prism GraphPad (San Diego, CA, USA).

For datasets with a limited number of animals (*N* < 5 per group), or when normality and homogeneity of variance could not be assumed, non-parametric tests were applied. Comparisons between two independent groups were performed using the Mann–Whitney U test (also referred to as the Wilcoxon rank-sum test). For analyses involving three or more independent groups, the Kruskal–Wallis test followed by Dunn’s post hoc test (with Holm correction for multiple comparisons) was used to identify pairwise differences. To assess the effect of two independent factors (e.g., *Treatment* and *Area*) and their potential interaction, we used the Scheirer–Ray–Hare test (SRH), a rank-based, non-parametric alternative to the two-way ANOVA. This method, implemented in both R and Python, extends the Friedman/Kruskal–Wallis approach to multifactorial designs, allowing analysis of main effects and interactions without assuming data normality.

Data are reported as median ± interquartile range (IQR), with individual animal values plotted for each group. All analyses were conducted considering the animal as the experimental unit, with slice-level data averaged per animal to avoid pseudoreplication.

## Results

### Differential effects of BoNT/A in severe and moderate SCI: combining rehabilitation for motor recovery and pain management

Our initial goal was to investigate whether intrathecal administration of BoNT/A during the onset of the chronic phase in paraplegic mice could restore motor function, as previously demonstrated when administered in the acute phase (12,13). Mice subjected to a severe thoracic spinal cord injury (T9–T11; see Supplementary Figure S1), as described in our earlier work (14), received a single intrathecal injection of either saline or BoNT/A (15 pg in 5 μL) at the lumbar level (L5). This injection site was chosen to minimize the risk of systemic toxin spread—particularly relevant in ischemic regions near the lesion—and to exploit the known retrograde transport capacity of BoNT/A to reach distant target (11,20,25).

However, in this preliminary study (Supplementary Figure S4), motor function assessed using the Basso Mouse Scale (BMS) score (17) showed that BoNT/A alone, administered during the chronic phase did not produce functional recovery, despite detectable central enzymatic activity, indicating that central modulation in isolation is insufficient in the presence of severe muscle atrophy.

These results led us to reexamine earlier studies (12,14,15) demonstrating that paraplegic animals experience pronounced muscle atrophy as early as day 7 post-trauma. Based on this, we hypothesized that the evaluation of BoNT/A’s effects, when administered in chronic phase, required a rehabilitative approach to mitigate muscle atrophy. To address this, we developed a hindlimb muscle electrostimulation (EMS) protocol using transcutaneous electrodes to deliver electrical stimulation from day 3 to day 15 post-trauma (protocol illustrated in Figure 1A and supplementary video S1). On day 15, animals were randomly divided into a control group (vehicle - saline) and a treatment group (BoNT/A), with EMS discontinued at this stage. Motor recovery was assessed throughout the EMS phase and after treatment (Figure 1B).

Animals undergoing EMS were classified as “severe” SCI cases, defined by complete hindlimb paralysis (BMS scores 0–3, where 0 indicates no movement and 9 indicates full motor recovery). Animals with BMS scores between 4 and 7, classified as “moderate” or “mild,” did not receive EMS but were included in the study to evaluate the toxin’s impact on neuropathic pain (NeP) during the chronic phase. Neuropathic assessments included responses to non-painful mechanical stimuli (dynamic aesthesiometer plantar test for allodynia), and painful thermal stimuli (plantar test for hyperalgesia). Given the variability in neuropathy onset between limbs, each paw was evaluated independently.

Figure 1C illustrates that severe SCI animals undergoing EMS rehabilitation showed no significant motor improvement up to day 15 post-lesion compared to non-EMS-treated SCI animals (all detailed statistical analysis are included in legend text). However, when BoNT/A or saline was spinally administered at day 15 and EMS was discontinued, only BoNT/A-treated mice exhibited gradual and significant motor recovery, reaching approximately 50% recovery in motor performance (supplementary video S2).

Regarding BoNT/A’s ability to mitigate NeP during the chronic phase, moderate and mild SCI animals showed significant prevention of allodynia compared to saline-treated SCI mice (Figure 1D). Additionally, while both groups developed hyperalgesia, BoNT/A treatment significantly reduced pain levels compared to saline-treated controls (Figure 1E).

### EMS in the Post-Acute Phase of SCI: A Rehabilitation Strategy to Prevent Muscle Atrophy and Facilitate BoNT/A Regenerative Action

Building on our previous findings (12, 14, 15), which demonstrated rapid muscle decline and atrophy as significant barriers to evaluating regenerative therapies in SCI, we considered clinical evidence suggesting that EMS in individuals with SCI can offer various benefits (26). These include increased muscle and bone mass, reduced fat mass, decreased spasticity, and improved functional mobility (27). To address hindlimb muscle atrophy in our severe SCI model, we implemented EMS prior to BoNT/A treatment to assess its potential in preventing or mitigating muscle loss.

SCI animals underwent the EMS protocol illustrated in Figure 1A and Supplementary Video 1. After 60 days, gastrocnemius (GA) muscles from different experimental groups (naïve, SCI without EMS, EMS+Saline, and EMS+BoNT/A) were collected and analysed (Figure 2).

**Figure 2.**
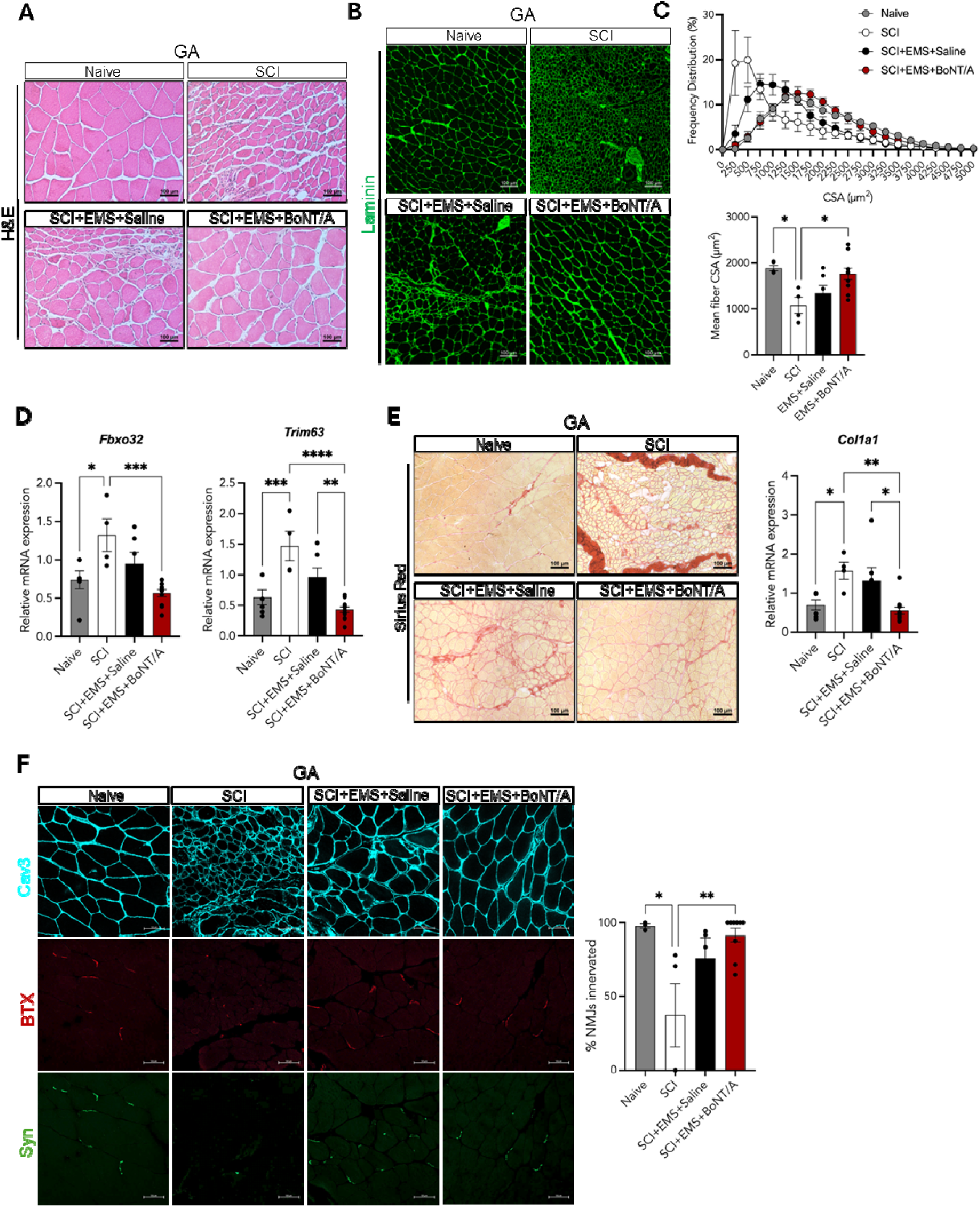
EMS and BoNT/A prevent muscle atrophy and preserve neuromuscular integrity in chronic SCI. **A)** Representative micrographs of gastrocnemius (GA) muscle sections from Naive, SCI (no EMS), EMS+Saline and EMS+BoNTA SCI mice stained with Haematoxylin-Eosin (H&E). Scale bars: 100 μm. **B)** Representative immunostaining for Laminin (green) in GA muscle sections from Naive, SCI (no EMS), EMS+Saline and EMS+BoNTA SCI mice. Scale bar,100 μm. **C)** Frequency distribution and mean of myofiber cross-sectional areas (CSA) measured on sections of GA muscle from Naive, SCI (no EMS ), EMS+Saline and EMS+BoNTA SCI mice; n ≥ 3, values represent mean ± SEM, Statistical significance was assessed by one-way ANOVA test: F (3, 21) = 5.752, p=0.0049. Post hoc tests for multiple comparisons between groups were performed by Tukey-Kramer test (*p < 0.05). **D**) Relative mRNA expression of Fbxo32 and Trim63 genes in GA muscle from Naive, SCI (no EMS), EMS+Saline and EMS+BoNTA SCI mice; n ≥ 4, values represent mean ± SEM. One-way ANOVA test: F (3, 23) = 7.690, p=0.0010; F (3, 24) = 14.23, p<0.0001, respectively. Post hoc tests for multiple comparisons between groups were performed with Tukey-Kramer test (*p < 0.05, **p < 0.01, ***p < 0.001, ****p < 0.0001). **E)** Representative micrographs of GA muscle sections from Naive, SCI (no EMS), EMS+Saline and EMS+BoNTA SCI mice stained with Sirius Red. Scale bars: 100 μm. The graph on the right shows the relative mRNA expression of Col1a1 gene in GA muscle from the same groups; n ≥ 4, values represent mean ± SEM One-way ANOVA test: F (3, 24) = 6.474, p=0.0023. Post hoc tests for multiple comparisons between groups were performed with Tukey-Kramer test (*p < 0.05, **p < 0.01). **F)** Representative immunostaining for caveolin-3 (Cav3) (cyan) and synaptophysin (Syn) (green) on GA muscle sections from Naive, SCI (no EMS), EMS+Saline and EMS+BoNTA SCI mice. NMJs are labelled with fluorescent alpha-bungarotoxin (BTX) (red). Scale bar, 50 μm. The graph on the right shows the percentage of innervated NMJs calculated by quantifying the Syn and BTX overlapping signals; n ≥ 3, values represent mean ± SEM. One-way ANOVA test: F (3, 16) = 5.315, p = 0.0098. Post hoc tests for multiple comparisons between groups were performed with Tukey-Kramer test (*p < 0.05, **p < 0.01)

Morphological analyses revealed that GA muscles from SCI-subjected mice exhibited a marked reduction in fibre size, quantified as a decrease in cross-sectional area (CSA) (Figure 2A-B-C). Specifically, we observed a significant shift toward a higher frequency of smaller fibres compared to naïve controls, alongside a significant reduction in mean fibre CSA. While EMS treatment alone showed a trend toward phenotypic improvement, a significant recovery of muscle morphology was only evident when EMS was combined with BoNT/A treatment (Figure 2A-B-C). We also examined the mRNA expression levels of FBXO32 (28) (also known as MAFbx/Atrogin-1) and TRIM63 (29) (MuRF1), two muscle-specific E3 ubiquitin ligases strongly induced during muscle atrophy. Consistent with the morphological data, both transcripts were significantly upregulated in the SCI group, reduced by EMS, and normalized in the EMS+BoNT/A group (Figure 2D). These results indicate that EMS applied during the post-acute phase of SCI exerts long-lasting protective effects against muscle atrophy and, when combined with the pro-regenerative activity of BoNT/A, promotes a substantial recovery of muscle homeostasis.

As previously reported, collagen deposition is a hallmark of disuse-associated muscle atrophy (12). Sirius Red staining revealed a marked accumulation of collagen in muscles from untreated SCI mice. This pathological alteration was clearly attenuated by EMS treatment and further improved by the combined EMS+BoNT/A protocol, indicating effective prevention of fibrotic deposition. Supporting this observation, the mRNA expression of collagen type I (Col1a1) was significantly upregulated in SCI muscles but prevented by EMS+BoNT/A treatment (Figure 2E).

Finally, we investigated whether the beneficial effects of the EMS+BoNT/A treatment were related to enhanced muscle innervation following spinal cord injury. As shown in Figure 2F, SCI induced a marked loss of muscle innervation, evidenced by the reduced colocalization of the postsynaptic marker bungarotoxin (BTX) with the presynaptic marker synaptophysin (Syn) at 60 days post-injury. EMS treatment partially preserved neuromuscular connectivity, whereas the combined EMS+BoNT/A protocol effectively maintained the integrity of neuron–muscle connections (Figure 2F).

Taken together, our findings highlight the potential of EMS, especially when combined with BoNT/A, as a promising rehabilitation strategy to counteract muscle atrophy, limit fibrotic changes, and preserve neuromuscular connectivity following severe SCI.

### Mechanisms of BoNT/A Neuroprotection: Mitigation of Excitotoxicity, Astocytosis and Reduction of Neuroinflammation

As previously demonstrated (12), and here confirmed (Supplementary Figure S5) BoNT/A has long-lasting enzymatic activity in the spinal cord acting on different cells type. We examined the presence of cleaved SNAP-25 in NeuN⁺ neurons and GFAP⁺ astrocytes within perilesional (T7 – T13) and epicentral regions (T9–T11) 60 days post-injury (Supplementary Figure S4). The yellow colocalization signal confirms persistent toxin activity in both neuronal and astrocytic populations. The detection of cleaved SNAP-25 at thoracic levels, distant from the lumbar injection site, supports the occurrence of retrograde migration of the toxin along the neuroaxis.

To investigate whether BoNT/A modulates astrocytic responses after SCI, we examined GFAP immunoreactivity and astrocyte distribution across the lesion epicenter (EPI), scar core (SCAR), and perilesional regions (PERI) at thoracic levels T7–T11, 60 days post-lesion (Figure 3A,B). Quantitative analysis revealed a region-specific effect of BoNT/A on astrocytic activation (Figure 3C). In EMS+Saline animals, GFAP staining showed the expected robust reactive profile at the lesion epicenter, with dense GFAP-positive processes and an elevated number of astrocytes. In contrast, EMS+BoNT/A treatment was associated with a marked reduction in both GFAP-positive area and astrocyte counts specifically within the epicentral tissue. This attenuation was not as evident in the surrounding perilesional zones, where GFAP coverage and astrocyte density remained comparable between groups, nor in the scar core, where high structural heterogeneity limited detectability of treatment-related differences.

**Figure 3.**
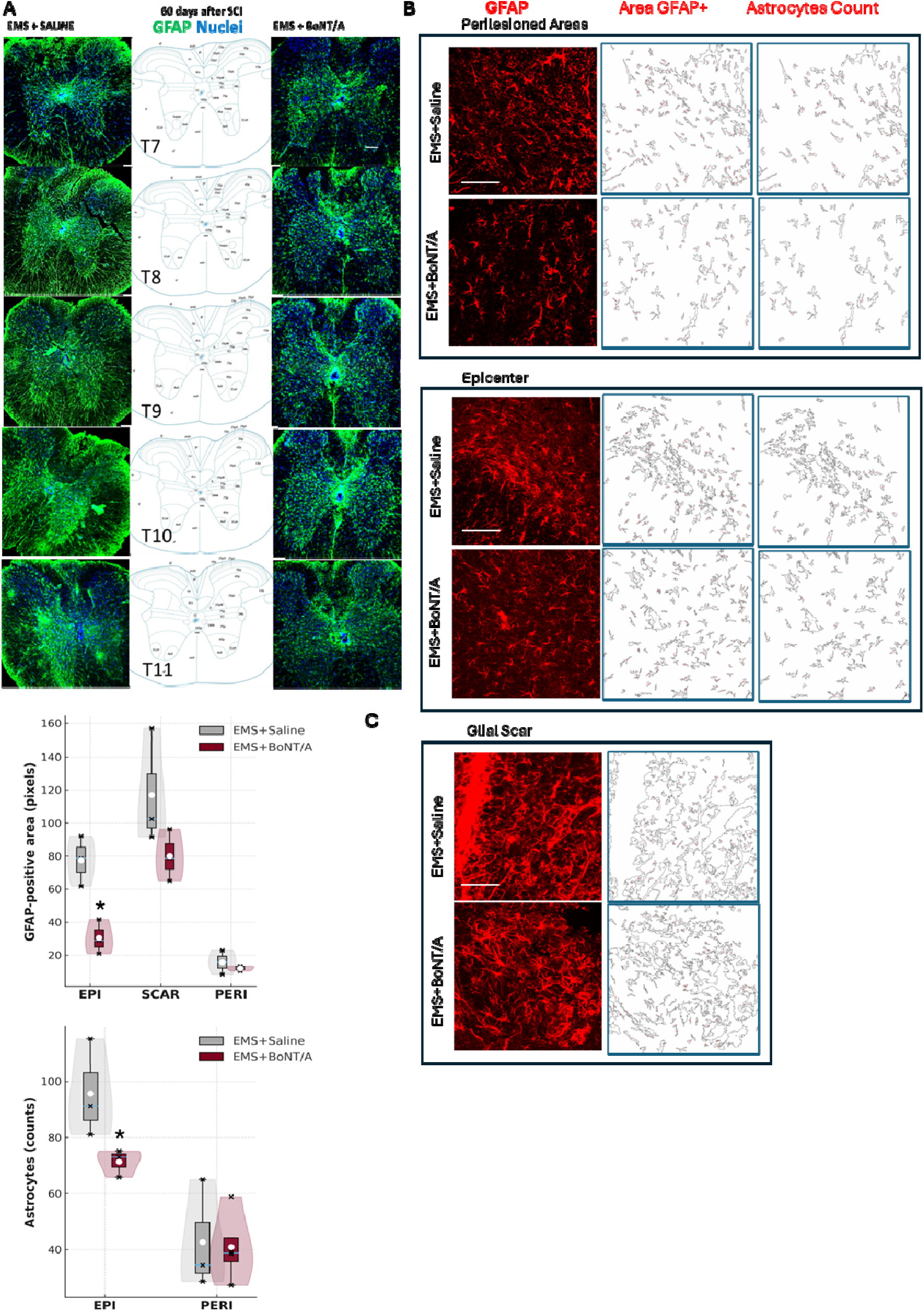
Modulation of astrocytic activation by BoNT/A at chronic stages after SCI. **(A)** Representative immunofluorescence images (10x, scale bar 100um) of GFAP-positive astrocytes (green) and nuclei (blue) across thoracic spinal cord levels (T7–T11), 60 days after SCI. Animals underwent EMS starting at day 3 post-injury and received either saline or BoNT/A at day 15 post-lesion. GFAP signal appears diffusely increased in EMS+saline mice, particularly at the lesion epicentre (T9–T10) Central columns show corresponding spinal cord atlas sections for anatomical reference. **(B–C)** Quantification of GFAP-positive area and astrocyte counts. GFAP images were converted to binary masks and processed using two segmentation thresholds to extract complementary structural information: a less restrictive threshold (“200 mask”) captured astrocyte somata together with GFAP-positive proximal processes, providing the measurement of GFAP-positive area (total GFAP+ pixels); a more restrictive threshold (“400 mask”) isolated astrocyte somata only, enabling automatic soma detection and astrocyte counting independent of processes or fragmented extensions. The masks shown next to the fluorescence panels illustrate these two segmentation outputs. A global comparison across treatment groups and regions (EPI, PERI, SCAR) using the Kruskal–Wallis test revealed a significant effect on GFAP-positive area: H_5_ = 15.76, p = 0.0076; Pairwise Mann–Whitney U tests showed: Epicentre (EPI): GFAP-positive area significantly reduced in EMS + BoNT/A mice vs EMS + Saline (p = 0.0495). Astrocyte number followed the same pattern, with a significant reduction in the epicentre of BoNT/A-treated animals (p = 0.0495). A Spearman correlation performed on combined EPI + PERI data demonstrated a strong positive association between astrocyte count and GFAP-positive area (ρ = 0.85, p < 0.001) indicating that slices with greater GFAP coverage also exhibited higher astrocyte soma density (N = 3–4 per group, ≥10 slices per animals; all slice-level values were averaged per animal, which was considered the experimental unit).

Across EPI and PERI regions, the relationship between GFAP-positive area and astrocyte number revealed a strong concordance (Spearman’s correlation analysis ρ = 0.85, *p* < 0.001), indicating that animals displaying more extensive GFAP immunoreactivity also showed higher astrocyte counts. Together, these results indicate that BoNT/A preferentially reduces astrocytic reactivity within the lesion epicenter, where glial activation is typically most pronounced. Effects in perilesional and scar territories appear more modest or variable, likely reflecting the intrinsic heterogeneity and structural complexity of these regions following chronic SCI.

To further characterize astrocytic morphology, we performed Sholl analysis on GFAP-positive cells from the different treatment groups (Figure 4A-C). The analysis of GFAP-positive astrocytes revealed a region- and treatment-dependent remodeling of astrocytic morphology. In the perilesioned area (Figure 4B), all injured groups showed increased intersections close to the soma, but SCI animals displayed the most pronounced perisomatic hypertrophy, with a higher number of proximal intersections than both EMS-treated groups. Within the EMS conditions, astrocytes from EMS+BoNT/A mice tended to show a slightly higher proximal intersection/radius profile but with a more compact arbor, as suggested by the reduced radial extension and process length. This pattern indicates that BoNT/A does not abolish proximal branching but rather favors dense, short-range processes around the soma. A similar organization emerged in the epicenter (Figure 4B, middle left), where tissue disruption is maximal. Here, SCI astrocytes again exhibited the largest and most extended arbors, whereas both EMS treatments reduced the radial spread of processes. Notably, EMS+BoNT/A astrocytes combined relatively high proximal intersections with a steeper decay of the Sholl curve and shorter maximum radius, resulting in the shortest and most compact arbors among the injured groups. The bar graphs summarizing intersection number and process length (Figure 4C, right) are consistent with this interpretation, highlighting a BoNT/A–associated confinement of astrocytic territorial extension both in the perilesioned rim and in the lesion core.

**Figure 4.**
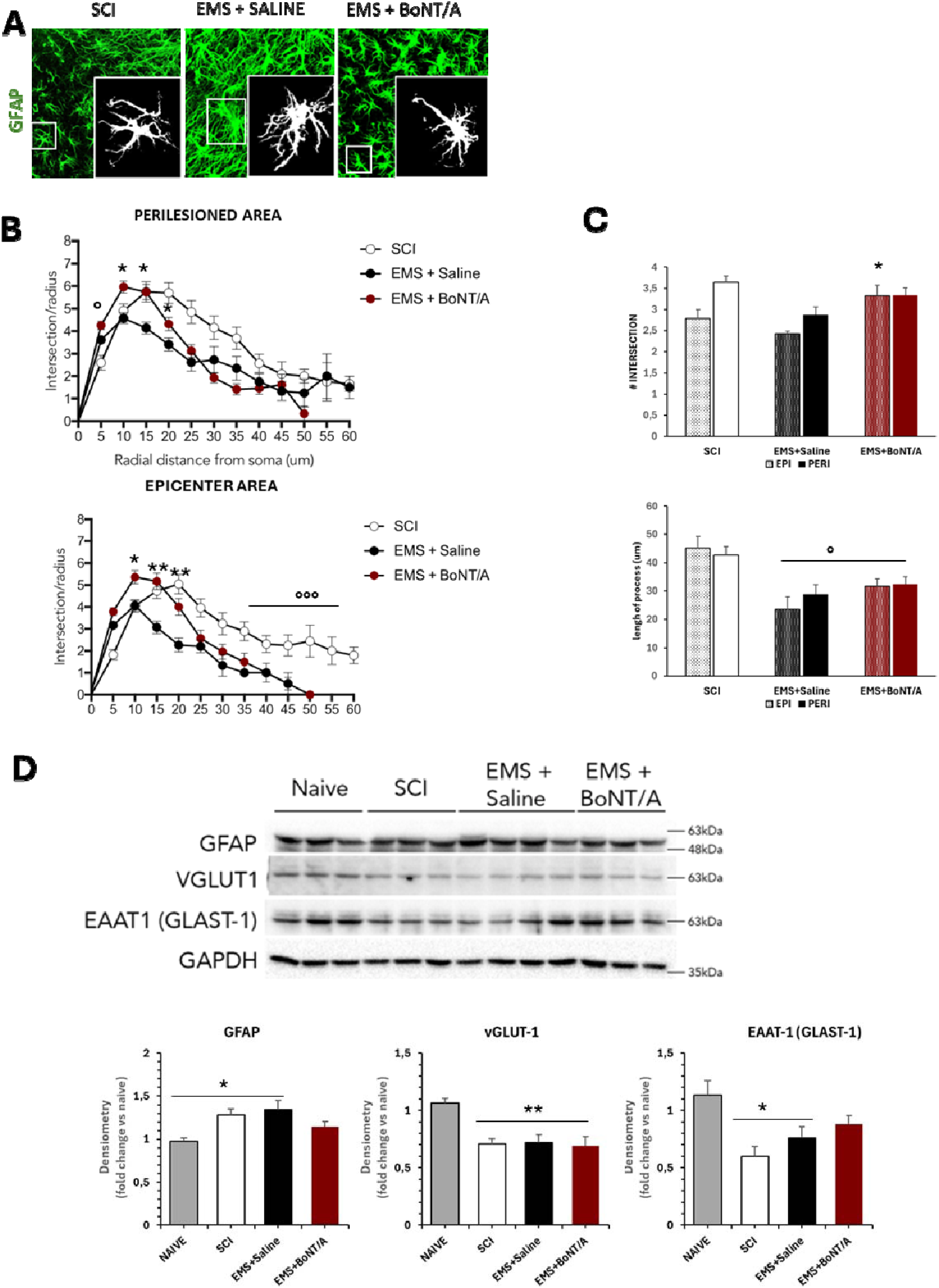
Astrocytic remodeling in the perilesioned and epicenter areas following SCI and EMS-based treatments. **A)** Representative extraction of GFAP+ cell from immunofluorescence images (40x zoom3) and corresponding binary reconstructions illustrate astrocyte morphology. Sholl profiles were generated by quantifying GFAP-positive intersections as a function of radial distance from the soma. **B)** Perilesioned Area (bottom graph). The Sholl analysis revealed significant differences restricted to the most proximal radii. A non-parametric Mann–Whitney test with FDR correction showed that SCI animals exhibited a significantly higher number of proximal intersections than both EMS+Saline and EMS+BoNT/A groups at radii 0–5 µm. A marginal difference was also detected for SCI vs EMS+Saline at 20–25 µm, whereas no significant differences emerged between EMS+Saline and EMS+BoNT/A at any radius after correction. Significance: *p < 0.05 vs EMS+Saline; ° p < 0.05 vs SCI. Epicenter Area (middle graph) Single-radius Mann–Whitney comparisons indicated significant differences between EMS+Saline vs EMS+BoNT/A at several proximal radii (5, 10, 15, 20 µm; all p < 0.001–0.0004). Kruskal–Wallis tests confirmed group effects on both the number of intersections (HL = 11.14, p = 0.0487) and the radial extent of processes (HL = 12.6, p = 0.0274). Moreover, quantitative Sholl-derived metrics confirmed that epilesional astrocytes in SCI animals display a markedly expanded arbor, with significantly greater AUC, maximal radius and estimated process length compared to both EMS-treated groups (Kruskal–Wallis p < 0.0001; Mann–Whitney SCI vs EMS+Saline p <0.0001; SCI vs EMS+BoNT/A p < 0.0001).**C)** Summary bar graphs (right). Average intersections (epicenter and perilesioned) and process length are shown for each group. Mann-Whitney post hoc tests revealed: *p < 0.05 vs EMS+Saline; ° p < 0.05 vs SCI. (n=3/4 group, slice/animal ≥5, cells/slice ≥3; all slice-level values were averaged per animal, which was considered the experimental unit) **D)** Western Blot Representative immunoblots for GFAP, vGLUT1, and EAAT1/GLAST-1 with GAPDH loading control. Densitometric quantification (fold change vs naïve) showed: GFAP: ANOVA *F*_3,21_ = 4.705, p = 0.0115; Tukey–Kramer: naïve vs saline *p* < 0.05, naïve vs SCI *p* < 0.05. vGLUT1: *F*_3,21_ = 7.68, p = 0.0012; Tukey–Kramer: naïve vs BoNT/A *p* < 0.001, naïve vs saline *p* < 0.001, naïve vs SCI *p* < 0.001. EAAT1/GLAST-1: *F*_3,21_= 5.102, p = 0.0083; naïve vs saline *p* < 0.05, naïve vs SCI *p* < 0.05 (N=6 naïve; 4 SCI; 8 EMS+ Saline; 5 EMS+BonT/A).

To further support the morphological findings, western blot analysis was performed (Figure 4D). GFAP levels were elevated in both SCI and EMS+Saline animals compared to naïve controls, indicating persistent astrocytic activation, whereas EMS+BoNT/A did not differ from baseline, suggesting attenuation of injury-associated GFAP upregulation. Similarly, vGLUT1, although classically associated with excitatory terminals (30), is also expressed at the astrocyte–synapse interface (31,32) and is increasingly recognized as a marker linked to glutamate handling and neuroinflammatory activity (32). All injured groups displayed a reduction in vGLUT1 compared to naïve tissue, with EMS+BoNT/A showing the most pronounced decrease, consistent with treatment-associated modulation of glutamatergic interfaces. EAAT1/GLAST-1 (33), a key astrocytic glutamate transporter, was also reduced after injury, particularly in SCI and EMS+Saline mice, whereas EMS+BoNT/A tended to preserve expression levels, pointing to a potential partial preservation of glutamate-buffering capacity. Second gel and ponceau staining are present in Supplementary Figure S9.

To confirm astrocytic–glutamatergic interactions, GFAP–vGLUT1 colocalization was evaluated, showing alteration after SCI and partially restored by BoNT/A (Supplementary Figure S6).

Taken together, these results indicate that BoNT/A modulates astrocytic reactivity, influencing structural complexity, glutamatergic interfaces, and associated molecular markers within the injured spinal cord.

Using high resolution CD11b/Iba1 imaging, microglial cells were categorized into five morphological phenotypes (Figure 5A) (34): - Resting (homeostatic): small soma with long, thin, highly branched processes that actively survey the surrounding environment; - Reactive: thicker and fewer processes, with a mildly enlarged soma, typical of inflammatory activation; - Amoeboid: rounded cells with minimal processes, associated with phagocytosis and strong immune activation; - Hyper-ramified: numerous long and thin processes, a morphology often linked to stress-related or transitional states; - Rod-shaped: elongated, polarized cells with very few processes, frequently aligned in chains and associated with chronic neuroinflammation or neurodegenerative conditions.

**Figure 5.**
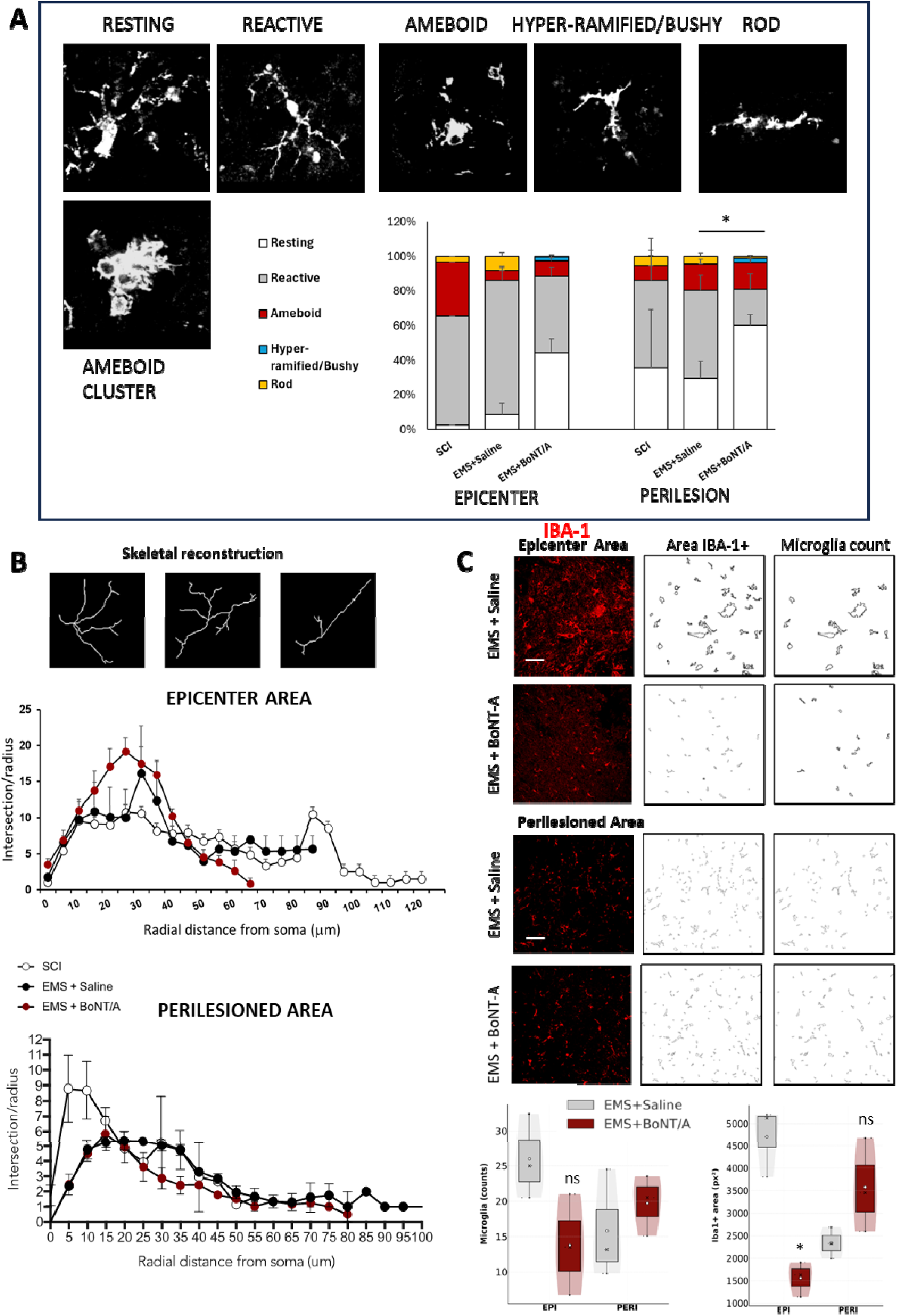
BoNT/A attenuates microglial activation and shifts microglial phenotype toward a resting state after SCI. **(A)** Microglial morphological phenotypes. Representative CD11b⁺/Iba1⁺ microglial morphologies identified in the epilesioned (EPI) and perilesioned (PERI) areas: Resting/Homeostatic, Reactive, Amoeboid, Hyper-ramified, and Rod-shaped. Quantification (count for section - animal-level mean of multiple slices; experimental unit = animal divided for EPI and PERI) revealed significant treatment-dependent differences in phenotype distribution in the perilesioned region. Mann–Whitney U tests: Resting %: BoNT/A > Saline *U = 0.000, Z = –1.964, p = 0.049;* Reactive %: BoNT/A < Saline; U = 0.000, Z = –1.964, p = 0.0495; Rod %: BoNT/A < Saline U = 0.000, Z = –1.964, p = 0.0495. Other phenotypes showed no significant differences. (N=3 animals/group, N=2 SCI; ≥ 7 sections/treatment). **(B)** Representative skeletonized reconstructions of Iba1⁺ microglia (up panel), illustrating the marked morphological variability across groups and the features quantified in the subsequent Sholl analysis. (Middle panel) Sholl analysis of microglia located within the lesion epicenter. Sholl curves represent average intersections per radius (animal-level means across multiple slices; experimental unit = animal). As expected for microglia residing at the epicenter, cells exhibited markedly higher branching densities and broader arborization profiles than in the perilesioned area (panel below). Global descriptors of arbor complexity (quantified as the number of intersections between microglial processes and concentric circles at increasing radial distances from the soma) computed on animal means did not reveal significant group differences: Area Under the Curve (AUC – total intersections): Kruskal–Wallis *H*L = 0.69, *p* = 0.707; Peak intersections: *H*L = 1.00, *p* = 0.607; Peak radius (distance of maximal branching from soma): *H*L = 2.88, *p* = 0.237. However, when the analysis was restricted to the 20–40 µm radial range, which encompasses the main branch-density peak, a robust group effect emerged: Kruskal–Wallis *H*L = 10.98, *p* = 0.0041 (radius-averaged intersections). Post-hoc Dunn’s comparisons indicated that microglia from EMS+BoNT/A-treated mice tended to display the highest intersection counts within this proximal domain, SCI microglia the lowest, and EMS+Saline an intermediate profile, consistent with a BoNT/A-driven enhancement of arbor complexity at the lesion epicenter. Sholl analysis of perilesioned microglia (down panel). Sholl curves reflect average intersections per radius (animal-level mean of multiple slices; experimental unit = animal). No significant group differences emerged in global arbor complexity: Area Under the Curve (AUC - total intersections): Kruskal–Wallis H_2_ = 5.14, p = 0.0766; Peak intersections: H_2_ = 4.03, df = 2, p = 0.1335. However, Peak radius (distance of maximal branching from soma) differed significantly among groups: Peak radius: H_2_ = 6.25, p = 0.0439. Post-hoc interpretation: SCI microglia reached their maximum branching closer to the soma (more compact, proximally hypertrophic profile), while both EMS-treated groups exhibited more distal branching. EMS+BoNT/A presented an intermediate profile between SCI and EMS+Saline. (N=3 animals/group, N=2 SCI; PERI ≥2 EPI ≥ 7 sections/treatment) **(C)** Microglial counts and Iba1⁺/CD11b⁺ area. Box/violin plots show microglial density and Iba1⁺ area in epilesioned (EPI) and perilesioned (PERI) regions. For Cell counts no significant differences were detected. For Iba1⁺ area EPI region: BoNT/A showed a significant reduction in microglial area compared with Saline U = 0.000, Z = –1.964, p = 0.0495. (N=3 animals/group, N=2 SCI; ≥ 7 sections/treatment). For PERI region, no significant difference, although BoNT/A tended to show larger areas. The larger Iba1⁺ area observed in the perilesioned BoNT/A group is compatible with its greater proportion of ramified microglia, which possess more extended arborisation. Conversely, saline tissue displayed more rod-shaped, reactive, and amoeboid morphologies, smaller, compact cell types that reduce the measured Iba1⁺ area despite being more activated.

Across chronic SCI tissue, multiple phenotypes coexisted within both epilesioned and perilesioned regions, though with different distributions (Figure 5A). In the perilesioned region, EMS+BoNT/A treatment produced a redistribution of morphology-based classifications toward a higher proportion of resting/homeostatic profiles, whereas EMS+Saline-treated animals exhibited a higher prevalence of reactive, ameboid, and rod-shaped microglia. In the epilesioned region, EMS+Saline animals also showed an enrichment of rod-shaped cells, while EMS+BoNT/A maintained a larger proportion of ramified morphologies (trend, p = 0.08).

Sholl profile quantification on microglia located at the lesion epicenter (Figure 5B) revealed a distinct pattern of branching complexity (quantified as the number of intersections between microglial processes and concentric circles at increasing radial distances from the soma) compared with the perilesioned region. At the epicenter, microglia from EMS+BoNT/A-treated mice displayed a broader and more distal branching profile, with increased process expansion within the proximal radial domain, whereas SCI microglia exhibited the most compact and centrally collapsed morphology. EMS+Saline microglia showed an intermediate pattern, indicating partial rescue of arbor structure following EMS alone. Overall, these profiles are consistent with a continuum ranging from the highly hypertrophic, process-retracted morphology typical of epicenter SCI microglia to a more extended, surveillance-like architecture restored by EMS+BoNT/A treatment.

Sholl profile quantification on perilesioned sections (Figure 5B) did not reveal statistically significant differences between groups. However, the overall pattern indicated that EMS+Saline-treated microglia displayed a more extended intersection/radius profile compared with BoNT/A, reflecting more compact arborisation in the toxin-treated group. This aligns with the coexistence of multiple phenotypes in chronic SCI tissue; hyper-ramified, reactive, and rod-like morphologies contribute to heterogeneous branching patterns that increase variability and dampen group-level statistical contrast.

Microglial cell counts (Figure 5C) did not differ significantly between groups in either region. In contrast, Iba1+ area was significantly reduced in the epilesioned region following EMS+BoNT/A treatment, indicating a lower burden of activated/reactive microglia compared with saline. In the perilesioned region, BoNT/A showed a trend toward increased Iba1+ area, consistent with the higher prevalence of ramified cells, which typically exhibit broader arborisation and therefore occupy a larger two-dimensional area despite being less reactive.

Importantly, EMS+Saline-treated tissue showed multiple clusters of ameboid and rod-shaped microglia, which are smaller, more compact phenotypes and therefore contribute to the reduced area despite being more reactive. The presence of these distinct and coexisting phenotypes explains why the morphometric outcomes (area, branching) exhibit higher variability in chronic SCI.

Together, these analyses depict a BoNT/A-mediated modulation of microglia toward less reactive states, particularly in the perilesioned region. While morphometric measures alone are influenced by the heterogeneity of phenotypes present in chronic tissue, the consistent pattern across morphology, area quantification, and Sholl profiles support the interpretation that BoNT/A is associated with a bias toward homeostatic-like ramified morphologies, contrasting with the predominance of reactive, ameboid, and rod-like microglia observed in saline controls.

For an integrated overview, the main results of EMS + BoNT/A on glial modulation are summarized in Table 2.

**Table 2.**
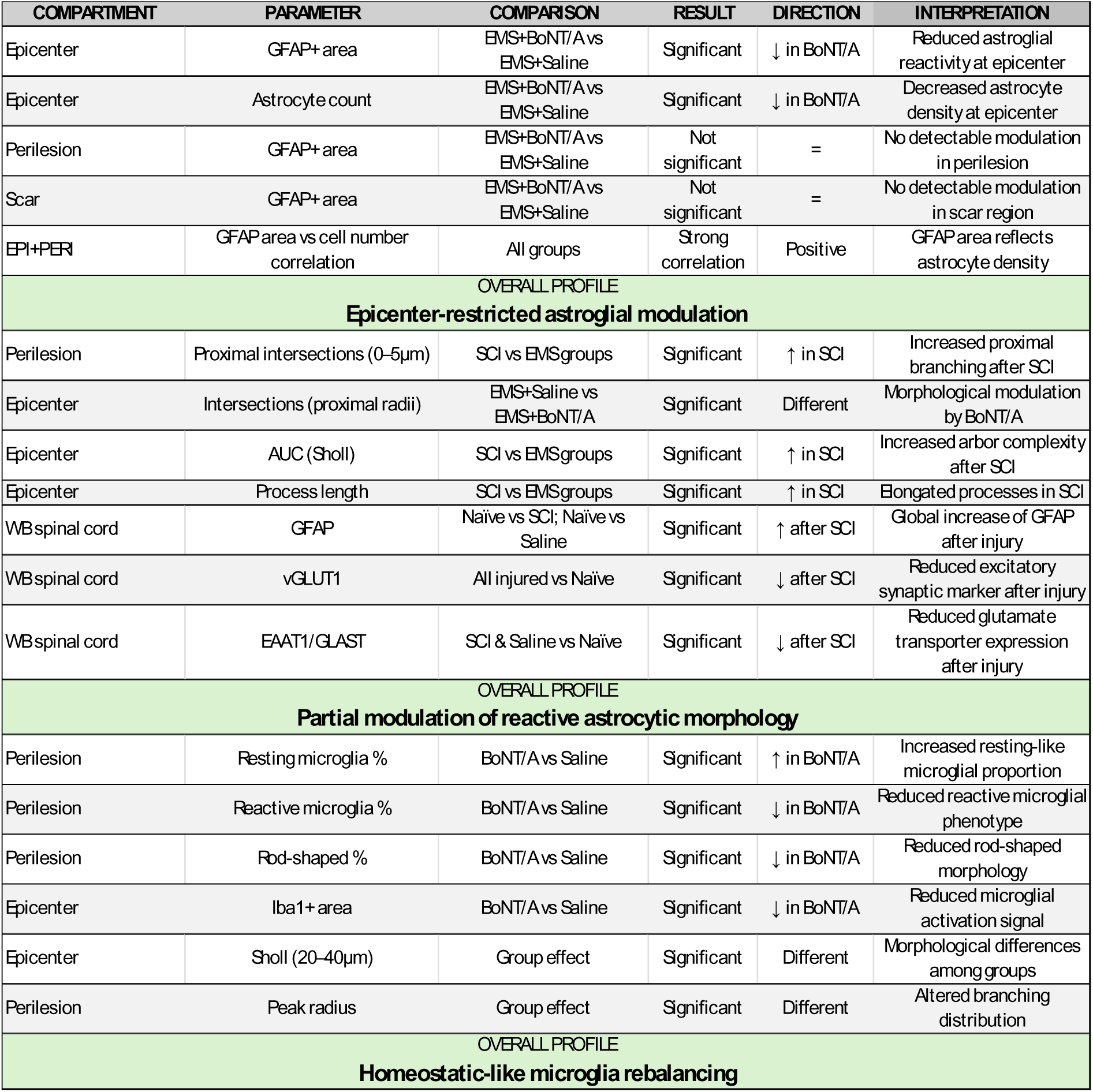
Summary of region-specific glial modulation induced by combined EMS + BoNT/A treatment in chronic SCI. The Table summarizes the principal glial endpoints (Figures 3-5) significantly affected by the combined EMS + BoNT/A treatment. The table reports compartment-specific analyses (epicenter, perilesion, scar), the statistical comparisons performed, and the direction of change. Directional arrows indicate variation relative to the specified comparator. The “Overall profile” sections provide a concise interpretation of the dominant trend emerging from each dataset without implying definitive cell-state transitions.

### BoNT/A reduces apoptotic cell death and promotes oligodendrocyte survival and differentiation after SCI

To assess whether astrocyte remodelling, glutamatergic modulation, and neuroinflammation reduction by BoNT/A contribute to neuroprotection, we evaluated apoptotic cell death in spinal cord tissue 60 days post-injury using TUNEL staining. Immunofluorescence analysis revealed a high density of TUNEL-positive nuclei (green) in both the SCI and EMS+vehicle groups, while markedly fewer apoptotic cells were observed in the EMS+BoNT/A group (Figure 6A). The spared parenchyma appeared more compact and better organized in BoNT/A-treated animals.

**Figure 6.**
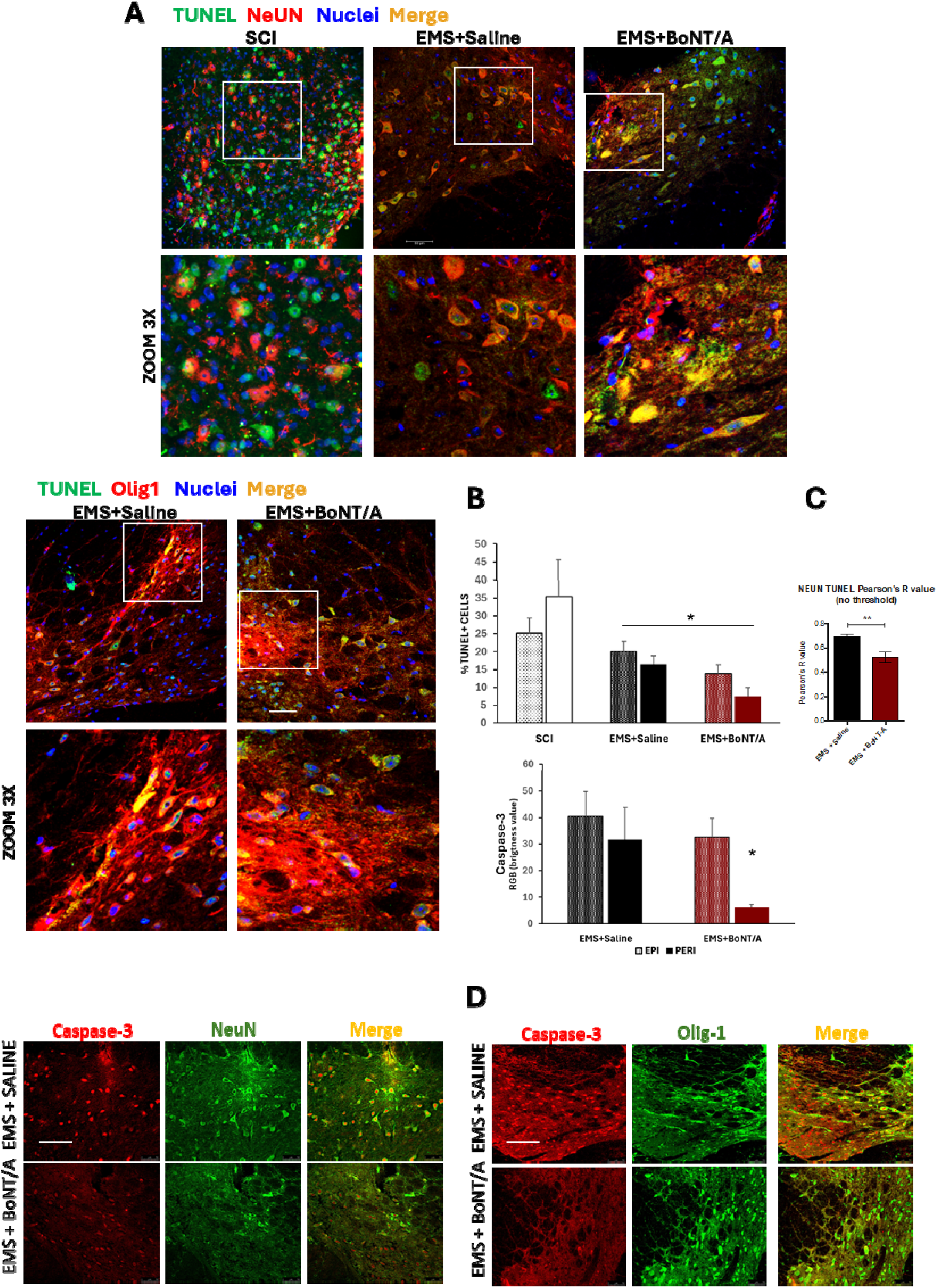
BoNT/A treatment reduces apoptotic cell death and preserves tissue integrity following SCI. **A)** Representative immunofluorescence images of transverse spinal cord sections stained for TUNEL (green), DAPI (blue), NeuN or Olig1 (red) and merged channels from SCI, EMS+Saline, and EMS+BoNT/A groups. Scale bar 50 μm. **B**) Quantitative analysis of apoptosis based on TUNEL staining. The percentage of apoptotic cells was calculated from the automated counts of TUNEL⁺ cells and total nuclei acquired for each slice. TUNEL staining was quantified in the lesion epicenter (EPI: T9–T11) and in perilesional regions (PERI: T7–T8 and T12–T13) in SCI, EMS+Saline, and EMS+BoNT/A–treated animals. Kruskal–Wallis analysis revealed a significant overall difference among groups (HL = 12.47, p = 0.0289), and post hoc pairwise comparisons (Mann–Whitney test) showed a significant reduction in apoptotic cells in EMS+BoNT/A compared with EMS+Saline in both epicenter and perilesional spinal regions (p < 0.05). **C)** Quantification of TUNEL-NeuN co-localization analysis. Graphs show a significant reduction in co-localization between neuronal marker NeuN and apoptotic TUNEL signal in BoNT/A-treated animals, assessed via Pearson’s correlation (no threshold, unpaired Student t-test: t_19_=2.882, **p<0.001). **D)** Representative images (40x) of Caspase-3 (red) immunoreactivity and its co-localization with NeuN or Olig1 (green). Scale bar 100μm. RGB intensity analysis (upper graph) did not show an overall significant difference among groups (Kruskal–Wallis test). However, post hoc pairwise comparisons revealed a significant reduction in signal intensity in the perilesional spinal regions of EMS+BoNT/A-treated animals compared with EMS+Saline mice (p<0.05). N = 3–4 animals/group, N=2 SCI with a minimum of 10 images analyzed for each animal, all slice-level values were averaged per animal, which was considered the experimental unit.

Quantification confirmed a general reduction in the proportion of apoptotic nuclei (TUNEL⁺/DAPI) in the EMS+BoNT/A group compared to both SCI and EMS+Saline groups (Figure 6B,C). This decrease was significant not only at the lesion epicenter (T9–T11) but also in more distant perilesional regions (T12–T13 and T7–T8). These findings support a pro-survival and neuroprotective action of BoNT/A within the injured spinal cord. To better illustrate how apoptotic cells distribute relative to the total number of nuclei across the tissue, individual values were plotted separately for the dorsal horn (DH) and ventral horn (VH), two regions that differ markedly in cellular density and composition, in the slice-wide distribution graph shown in Figure S8.

To determine whether this effect reflects preferential protection of specific cell types, we assessed the degree of co-localization between neuronal nuclei (NeuN⁺) and TUNEL staining. Pearson’s correlation analyses revealed significantly reduced co-localization in the EMS+BoNT/A group compared to EMS+Saline, suggesting that BoNT/A may preferentially protect neurons from apoptosis (Figure 6C). Additional analyses using thresholded Manders’ coefficients showed consistent trends but did not reach statistical significance (see Supplementary Figure S8).

We next examined the expression of the executioner protease Caspase-3 in the spinal cord (Figure 6D) parenchyma. Immunofluorescence analysis didn’t reveal a qualitative reduction of Caspase-3 signal for all analysed sections. However, a significant reduction in Caspase-3 signal intensity in the perilesioned spinal regions of EMS+BoNT/A–treated animals compared with EMS+Saline mice was appreciable. These findings indicate that BoNT/A reduces caspase-related apoptotic signaling specifically in areas surrounding the lesion, consistent with its broader neuroprotective profile.

To investigate whether the observed neuroprotection could involve restoration of excitatory/inhibitory receptor balance, we analyzed the expression and distribution of the NMDA receptor subunit ε2 and the GABA-A receptor subunit α2 in the spinal cord 60 days after injury (Figure 7A–D).

**Figure 7.**
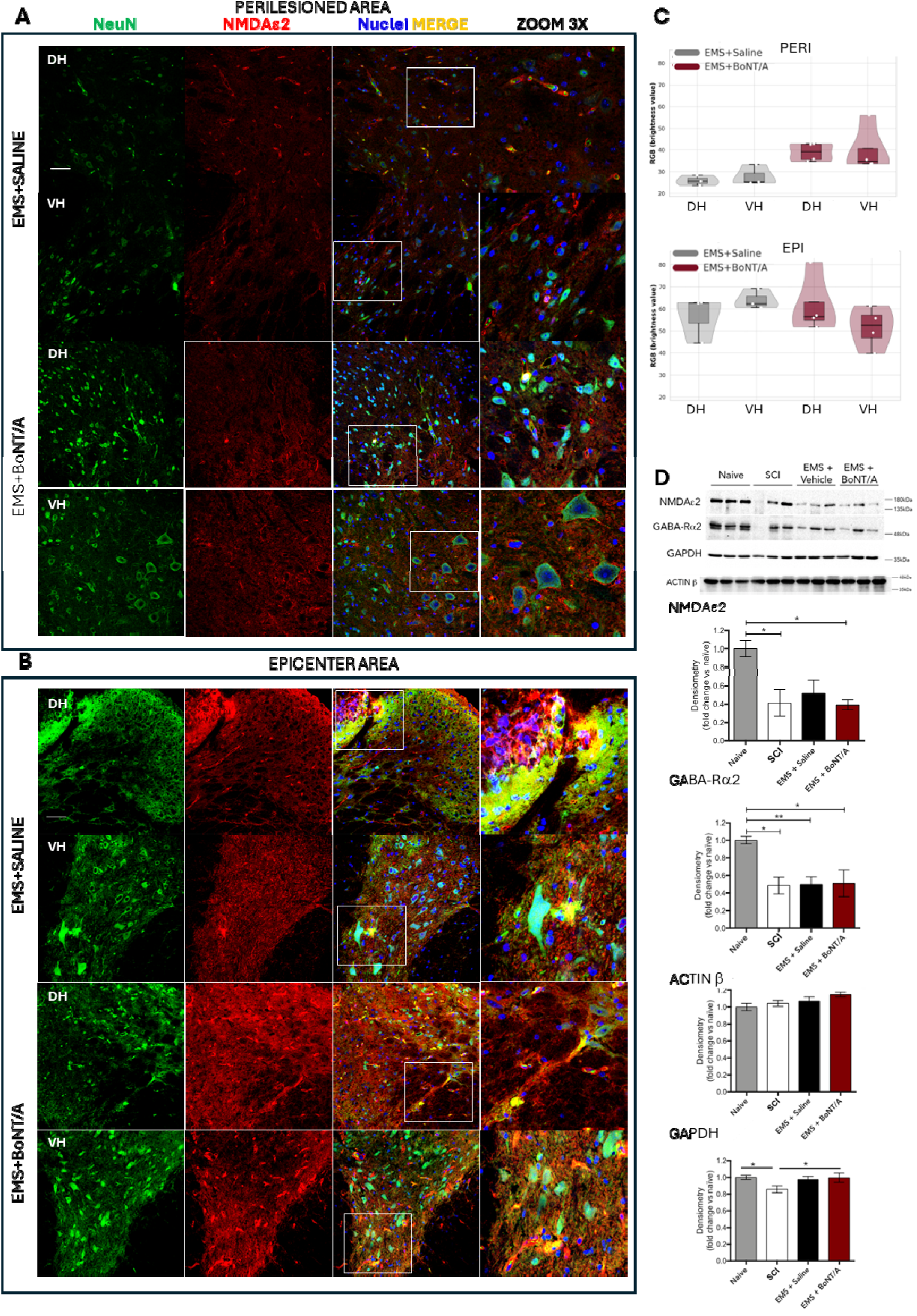
Expression of NMDA and GABA-A receptor subunits in the spinal cord 60 days after SCI. (A–B) Localization and distribution of NMDA receptor (NMDA ε2) immunoreactivity in the dorsal (DH) and ventral horns (VH) of the spinal cord. **(A)** Representative confocal images of perilesional regions (rostral–caudal, within 2–3 mm from the impact zone; T7–T13 segments). **(B)** Representative images of the epicentral area (T9–T11), corresponding to the region directly affected by the trauma. Sections were co-stained for NeuN (neuronal marker, green), NMDA ε2 (red), and nuclei (DAPI, blue). Images were acquired at 40× magnification; scale bar = 50 µm. **(C)** Quantitative analysis of NMDA ε2 fluorescence intensity in DH and VH across epicentral and perilesional regions. Each point represents one animal (*N* = 3–4 per group). All slice-level values (two per animal for each area and two DH/VH measurements per slice) were averaged per animal, which was considered the experimental unit. Boxplots display median ± IQR, and violin plots illustrate data distribution. Kruskal–Wallis analysis indicated a significant overall difference among the four groups for both DH (*p* = 0.0117) and VH (*p* = 0.0139); however, no pairwise comparison remained significant after Holm correction. Scheirer–Ray–Hare testing revealed a main effect of Area (EPI vs PERI) on both DH (*p* = 0.0017) and VH (*p* = 0.0060), with no significant main effect of Treatment and no significant interaction (trend for VH, *p* = 0.080). **(D)** Western blot analysis (*N* = 6–7 per group) of excitatory (NMDA ε2) and inhibitory (GABA-A α2) receptor subunits in spinal cord lysates collected 60 days after injury. Representative blots (left) show protein levels across groups. Quantification (right) was performed on all available samples (Naïve *n* = 6; SCI *n* = 4; EMS+Saline *n* = 7; EMS+BoNT/A *n* = 4) and expressed as fold-change relative to the naïve mean. Both receptors were significantly modulated in injured groups versus naïve controls, with no major differences among SCI, EMS+Saline, and EMS+BoNT/A animals (one-way ANOVA: NMDA ε2 *F*_3,21_ = 5.48, *p* = 0.0061; GABA-A α2 *F*_3,21_ = 2.34, *p* = 0.0037). GAPDH and β-actin levels also varied across groups (unpaired t-tests: GAPDH naïve vs SCI *t*LL = 2.92, *p* = 0.0154; Actin BoNT/A vs Naïve *t*LL = 2.91, *p* = 0.0061). Protein loading was verified by Ponceau staining (see Supplementary Figure S5).

Immunofluorescence analysis of NMDA receptor (NMDAε2) distribution (Figure 7A–C) revealed the spatial organization of NMDA ε2 labeling within the dorsal (DH) and ventral (VH) horns of both epicentral and perilesional regions. Qualitatively, NMDAε2 immunoreactivity appeared more intense in perilesional areas compared to the lesion core, a pattern observed in both EMS+Saline and EMS+BoNT/A groups. A trend toward higher NMDA signal in the perilesional region of BoNT/A-treated mice was detected, although not supported by post hoc statistics. The same pattern of distribution has been observed in GABA-A receptor subunit α2 (GABA-Rα2) but without evident signs of differences between two groups analyzed (Supplementary Figure S9).

Western blot analysis (Figure 7D) confirmed that both NMDA ε2 and GABA-R α2 receptor subunits were significantly downregulated in all injured groups compared to naïve controls, with no differences among SCI, EMS+Saline, and EMS+BoNT/A animals, thereby confirming a persistent synaptic imbalance that is not corrected by BoNT/A treatment.

Interestingly, we also observed treatment- and injury-dependent modulation of β-actin and GAPDH, two commonly used housekeeping proteins. Specifically, GAPDH levels were reduced in SCI tissue, while β-actin was increased in the EMS+BoNT/A group. These changes may reflect broader cytoskeletal and metabolic remodelling processes in response to injury and treatment. Notably, normalization of receptor signals to GAPDH abolished statistical significance (data not shown), highlighting the potential instability of conventional loading controls in lesioned tissue. Ponceau staining confirmed consistent protein loading across lanes (see Supplementary Figure S9).

To further characterize the cytoprotective effects of BoNT/A, we examined oligodendroglia survival and myelin protein expression in the injured spinal cord 60 days post-injury. Immunofluorescence for Olig-1 revealed (Figure 8A) a higher density and intensity of Olig1⁺ cells in EMS+BoNT/A-treated animals compared to EMS+Saline and the evaluation of fluorescence brightness of Olig1 expression in the BoNT/A group reveal an increasing trend, although not significant both in epicenter or perilesioned areas.

**Figure 8.**
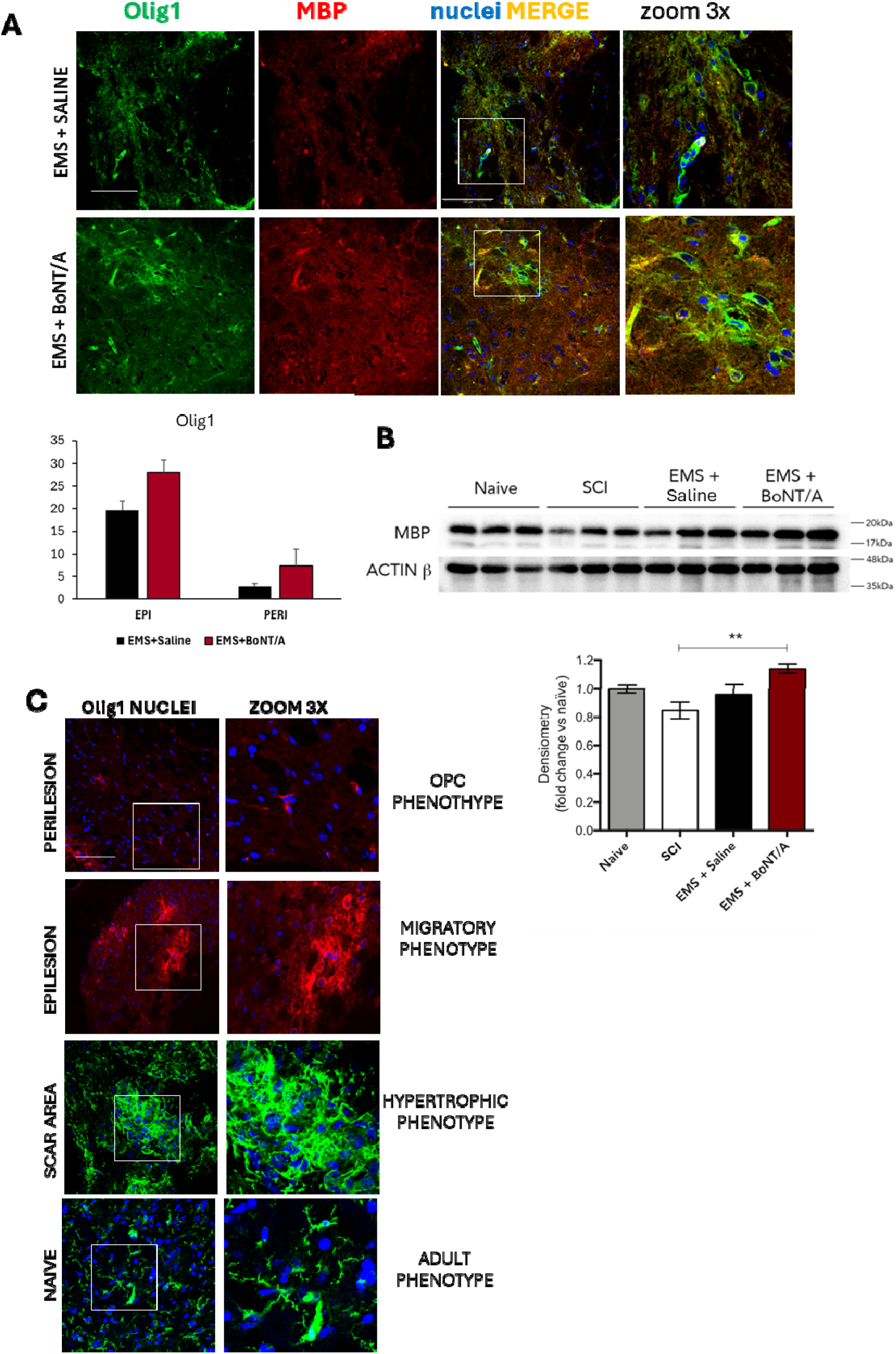
BoNT/A modulates Olig1 expression and enhances myelin protein expression in the chronic phase of SCI. **A)** Representative immunofluorescence images (scale bar 100 μm) of spinal cord sections stained for Olig1 (green), MBP (red), and DAPI (blue) from EMS+Saline and EMS+BoNT/A groups, acquired 60 days after injury. Merged images and 3× zoom-in panels illustrate Olig1L cells and enhanced MBP signal in BoNT/A-treated animals. Quantification of fluorescence brightness values, separately evaluated in the epicenter (EPI) or perilesioned (PERI) areas shows a trend, although not significant, in increased Olig1 expression in the EMS+BoNT/A group compared to EMS+Saline. N=2/3 animals/treatment, Saline: 12-40 slices, BoNT/A: 10-36 slices; all slice-level values were averaged per animal, which was considered the experimental unit). **B)** Representative Western blot of MBP in spinal cord lysates from all experimental groups (n = 3 per group shown in blot). Densitometric analysis of MBP levels (normalized to the average of naïve controls) is shown in the bar graph, including all samples: naïve (n = 6), SCI (n = 4), EMS+Saline (n = 7), EMS+BoNT/A (n = 4). One-way ANOVA revealed a significant group effect (F_3,21_ = 5.153; p = 0.0079), with Tukey-Kramer post hoc test indicating a significant increase **of** MBP in the EMS+BoNT/A group compared to SCI (p < 0.01) .The modulation of MBP expression is not statistically significant when normalized to ACTIN-β, due to variability in housekeeping gene levels across groups (as shown in figure 6). Protein loading was assessed via Ponceau staining (Supplementary Figure S9), and a replicate Western blot is shown in Supplementary Figure S9. **C)** Distinct oligodendrocyte phenotypes expressing Olig1 were similarly detected in both Saline- and BoNT/A+EMS-treated mice. Confocal images (40x, scale bar 100μm) representing Olig1 staining in EPI-, PERI-lesion area, and in scar and naïve tissue in which is appreciable a change in phenotypic profile. The RGB values shown in panel A) reflect phenotype-specific changes in oligodendrocytes driven by the injury and by the anatomical region affected. For this reason, areas corresponding to the glial scar were excluded from the RGB-based analysis.

To determine whether BoNT/A also influenced oligodendrocyte maturation and myelin production, we analyzed myelin basic protein (MBP) expression. Western blot results showed a significant increase in MBP levels in EMS+BoNT/A animals compared to all other groups, including naïve, SCI, and EMS+vehicle (Figure 8B). Quantitative densitometry confirmed this effect, indicating enhanced myelin protein synthesis in response to BoNT/A treatment. Of interest, morphological observation (Figure 8C) of spinal cord sections collected from distal (perilesion) and proximal (epilesion) regions, as well as from the glial scar, in both EMS+Saline- and EMS+BoNT/A-treated mice revealed a marked shift in Olig1 localization and oligodendrocytes phenotype, consistent with previous reports (35–37).

To verify whether BoNT/A exerts a direct effect on the oligodendroglia lineage (Figure 9), we exposed cultured oligodendrocyte precursor cells (OPCs) to 10 pM BoNT/A under either proliferative (NO DIFF) or differentiating (DIFF) conditions. Cell counts at 24- and 48-hours post-treatment showed no significant differences in OPC survival across groups (Figure 9A), indicating that BoNT/A does not compromise cell viability.

**Figure 9.**
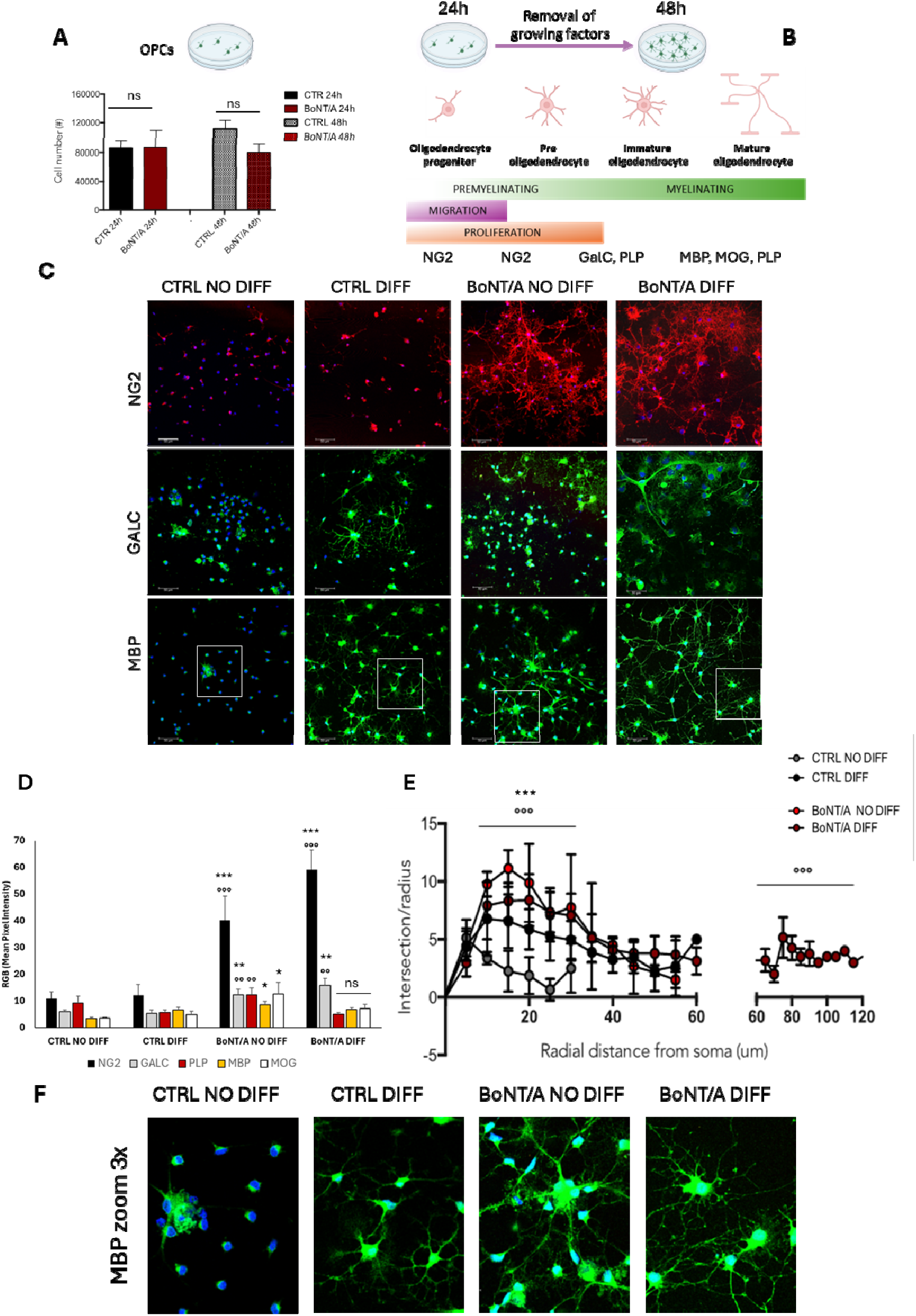
BoNT/A enhances oligodendroglia differentiation and morphological complexity in vitro. (**A**) Timeline of the experimental protocol: oligodendrocyte progenitor cells (OPCs) were cultured under proliferative conditions for 24 h and then switched to differentiating conditions (48 h), in the presence or absence of BoNT/A. **(B)** Schematic of the oligodendrocyte lineage progression: OPCs differentiate into pre-oligodendrocytes, immature oligodendrocytes, and finally mature myelinating cells. Marker expression during this process includes NG2 (Neuron-glial antigen 2, OPC marker), GalC (Galactocerebroside, immature OL marker), PLP (Proteolipid protein), MBP (Myelin basic protein), and MOG (Myelin oligodendrocyte glycoprotein) as markers of mature myelinating oligodendrocytes.**(C)** Representative confocal images (40×, scale bar = 50 μm) of NG2 (red), GalC (green), and MBP (green) staining in control and BoNT/A-treated cultures under differentiating (DIFF) and non-differentiating (NO DIFF) conditions. Nuclei are counterstained with DAPI (blue). (**D)** Quantitative analysis of immunofluorescence intensity (mean RGB values – N=3 group; 6-12 images evaluation/group) revealed the following: NG2: ANOVA F_3,20_= 13.834, p < 0.0001. Tukey-Kramer post-hoc: BoNT/A NO DIFF vs CTRL NO DIFF p < 0.005; BoNT/A NO DIFF vs CTRL DIFF p < 0.005; BoNT/A DIFF vs CTRL NO DIFF p < 0.0001; BoNT/A DIFF vs CTRL DIFF p < 0.0001; GalC: ANOVA F_3,34_= 8.037, p = 0.0004. Tukey-Kramer post-hoc BoNT/A NO DIFF vs CTRL NO DIFF p < 0.001; BoNT/A NO DIFF vs CTRL DIFF *p < 0.05*; BoNT/A DIFF vs CTRL NO DIFF *p < 0.*0005; BoNT/A DIFF vs CTRL DIFF *p < 0.05*; PLP: ANOVA F_3,16_ = 4.089, p = 0.0248. Tukey-Kramer post-hoc BoNT/A NO DIFF vs CTRL DIFF *p < 0.05*; MOG: ANOVA _F3,20_ = 2.59, p = 0.081. Tukey-Kramer post-hoc BoNT/A NO DIFF vs CTRL NO DIFF *p < 0.05*; MBP: ANOVA F_3,21_= 4.145, p = 0.0187. Tukey-Kramer post-hoc BoNT/A NO DIFF vs CTRL NO DIFF *p < 0.05*. **(E)** Sholl morphometric analysis of MBP⁺ cells performed using the Neuroanatomy plugin in FIJI/ImageJ. Repeated Measure ANOVA shows main effect for Treatment (F_3,193_=98,53 p<0.0001) for Radial distance (F_12,192_=60,08 p<0.0001) and interaction Treatment x Radial distance (F_36,192_=5,53 p<0.0001) Tukey-Kramer post-hoc test evidenced significant differences in number of intersections per radius: BoNT/A DIFF cells showed significantly increased arborization compared to CTRL NO DIFF and CTRL DIFF up to 20 µm from the soma (*p < 0.001*). BoNT/A NO DIFF vs CTRL NO DIFF at the same radial distance (*p < 0.*0001) and for total process length: Both BoNT/A DIFF and BoNT/A NO DIFF groups showed significantly longer total process length compared to CTRL NO DIFF (*p < 0.*0001). **(F)** High magnification images (zoom 3×) of MBP immunofluorescence (showed in panel C – area in the square) highlighting the increased complexity and arborization in BoNT/A-treated cells.

Morphological analysis using immunofluorescence revealed distinct changes in marker expression across the maturation stages of the oligodendroglia lineage (Figure 9C). Under control differentiating conditions, a partial increase, in GalC⁺ (galactocerebroside C) and MBP⁺ cells, was observed, consistent with normal progression toward immature and mature oligodendrocytes. However, BoNT/A treatment, both in proliferative and differentiating conditions, led to a significant upregulation of GalC, MBP, and MOG (myelin oligodendrocyte glycoprotein) immunoreactivity, suggesting accelerated differentiation and myelin acquisition. RGB quantification confirmed these effects. Expression’s evaluation confirmed that BoNT/A promotes the differentiation of oligodendrocyte lineage cells.

In particular, GalC, a marker of immature oligodendrocytes, was strongly upregulated in both BoNT/A-treated groups, under proliferative (NO DIFF) and differentiating (DIFF) conditions, indicating a robust shift toward a premyelinating phenotype. Expression of MBP was also elevated following BoNT/A treatment, even in the absence of differentiation stimuli, suggesting an early initiation of myelin-related programs. MOG (myelin oligodendrocyte glycoprotein) levels showed a similar trend, further supporting the progression toward a mature, myelin-producing oligodendrocyte phenotype. Notably, PLP (proteolipid protein), another marker of mature oligodendrocytes, was selectively increased in BoNT/A-treated cells maintained in proliferative conditions, reinforcing the ability of BoNT/A to drive maturation independently of extrinsic differentiation cues (representative images of PLP- and MOG-positive OPC staining are provided in Figure 10). In contrast, NG2 (neuron-glial antigen 2), a marker of oligodendrocyte precursor cells, was significantly downregulated in all BoNT/A-treated groups, consistent with the exit from the progenitor state and commitment to the differentiation pathway.

**Figure 10.**
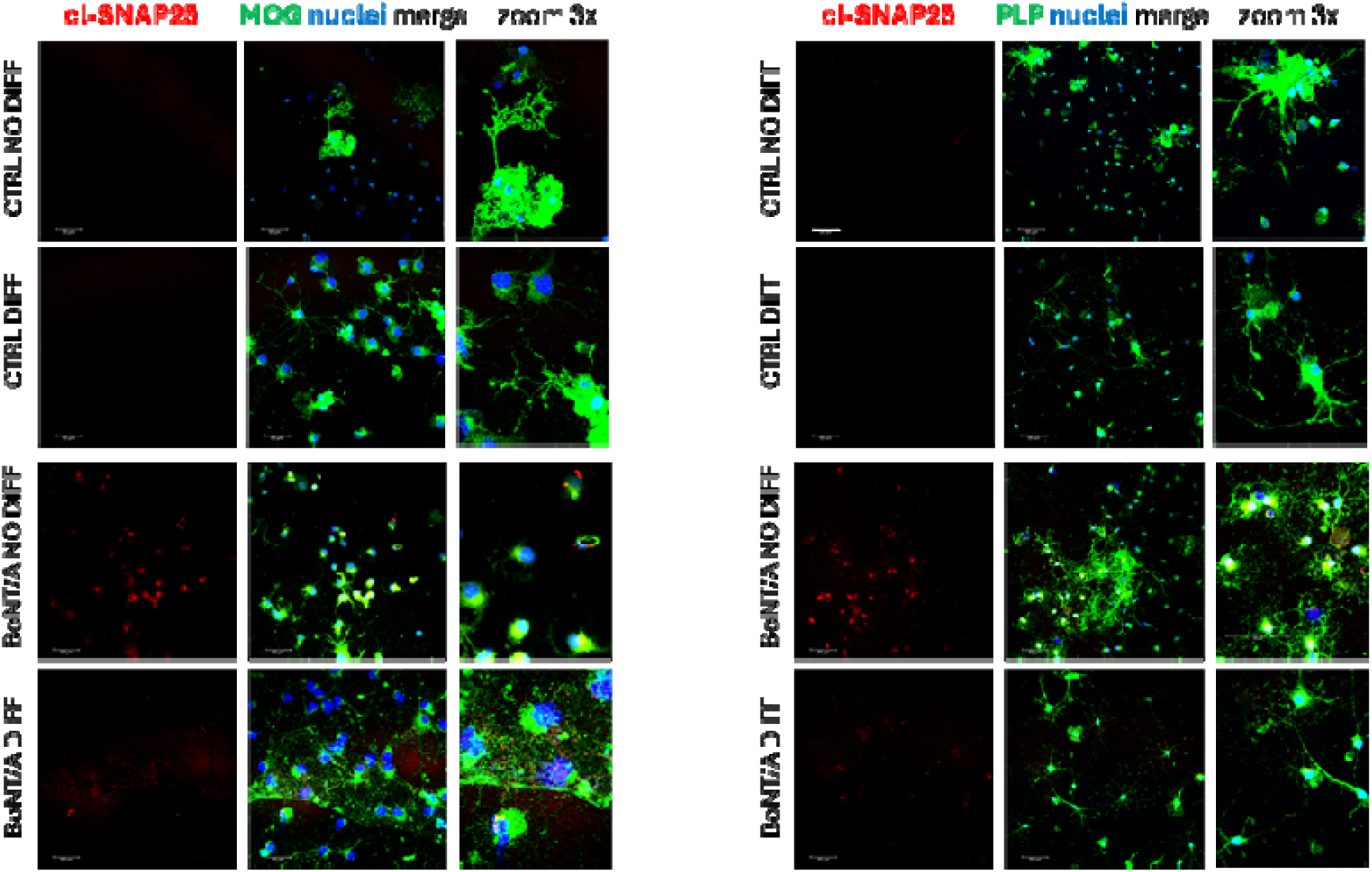
BoNT/A internalization in oligodendroglial cells demonstrated by colocalization with cl-SNAP25. Representative confocal images (40×, scale bar = 50 μm) of primary oligodendrocyte cultures stained with antibodies against cl-SNAP25, (red), the oligodendrocyte maturation markers MOG (Myelin Oligodendrocyte Glycoprotein, green, left panel) or PLP (Proteolipid Protein, green, right panel), and nuclear counterstain (DAPI, blue). BoNT/A uptake and functional enzymatic activity were evaluated by immunolabeling for cl-SNAP25, a specific marker of BoNT/A-mediated SNAP25 cleavage. Images show partial or complete colocalization between cl-SNAP25 and MOG⁺ or PLP⁺ cells

To assess whether BoNT/A enhances morphological complexity, we performed Sholl analysis on MBP⁺ cells (Figure 9E). BoNT/A-treated cells exhibited a significantly higher number of intersections at distances up to 20 μm from the soma compared to CTRL NO DIFF and CTRL DIFF. In addition, both BoNT/A NO DIFF and DIFF groups showed significantly increased total process length vs CTRL NO DIFF, further supporting the toxin’s effect on promoting arborization and maturation.

High-magnification images (Figure 9F) confirmed the increased complexity and branching of MBP⁺ cells in BoNT/A-treated cultures, even under non-differentiating conditions.

To confirm that BoNT/A is internalized and enzymatically active in oligodendrocytes, we performed triple immunofluorescence for cleaved SNAP25 (cl-SNAP25), MOG or PLP, and DAPI (Figure 10). Confocal imaging revealed clear colocalization of cl-SNAP25 signal within MOG⁺ and PLP⁺ cells, indicating that BoNT/A enters oligodendroglia cells and cleaves SNAP25, its canonical substrate. This result provides direct evidence of BoNT/A internalization and activity in cells of the oligodendrocyte lineage.

Taken together, these in vitro findings demonstrate that BoNT/A directly promotes the differentiation of OPCs toward mature myelinating oligodendrocytes.

For an integrated overview, the main results of EMS + BoNT/A on neuroprotection are summarized in Table 3.

**Table 3.** Summary of neuroprotection and oligodendroglial responses induced by combined EMS + BoNT/A treatment in chronic SCI. Table summarizes the principal apoptosis-related and oligodendroglial endpoints (Figures 6, 8, 9) significantly modulated by EMS + BoNT/A treatment. The table reports the anatomical compartment analyzed, the statistical comparison performed, and the direction of change relative to the specified comparator. “Overall profile” statements synthesize the dominant trend emerging from each dataset and reflect modulation of apoptosis-associated markers and oligodendroglial maturation without implying direct quantification of net cell survival or complete remyelination.

| COMPARTMENT | PARAMETER | COMPARISON | RESULT | DIRECTION | INTERPRETATION |
| --- | --- | --- | --- | --- | --- |
| Epicenter | TUNEL+ cells | EMS+BoNT/A vs EMS+Saline | Significant | ↓ in BoNT/A | Reduced apoptotic signal |
| Perilesion | TUNEL+ cells | EMS+BoNT/A vs EMS+Saline | Significant | ↓ in BoNT/A | Reduced apoptotic signal |
| Epicenter | NeuN+/TUNEL+ colocalization | EMS+BoNT/A vs EMS+Saline | Significant | ↓ in BoNT/A | Reduced neuronal apoptotic involvement |
| Perilesion | Caspase-3 signal | EMS+BoNT/A vs EMS+Saline | Significant | ↓ in BoNT/A | Reduced apoptotic marker expression |
| OVERALL PROFILE |  |  |  |  |  |
| Reduction of apoptosis-associated markers |  |  |  |  |  |
| Epicenter | Olig1+ density | EMS+BoNT/A vs EMS+Saline | Significant | ↑ in BoNT/A | Increased oligodendroglial presence |
| Perilesion | Olig1+ density | EMS+BoNT/A vs EMS+Saline | Significant | ↑ in BoNT/A | Increased oligodendroglial presence |
| WB spinal cord | MBP | EMS+BoNT/A vs EMS+Saline | Significant* | ↑ in BoNT/A | Increased myelin-associated protein |
| Epicenter | Oligodendroglial maturation markers | EMS+BoNT/A vs EMS+Saline | Significant | ↑ in BoNT/A | Enhanced maturation profile |
| In vitro OPC differentiation | Differentiated phenotype | BoNT/A vs Control | Significant | ↑ in BoNT/A | Promotion of oligodendroglial differentiation |
| OVERALL PROFILE |  |  |  |  |  |
| Pro-remyelinating oligodendroglial response |  |  |  |  |  |

## Discussion

Chronic SCI represents one of the most pressing and unresolved challenges in neurorehabilitation. Although acute SCI has been the focus of most clinical and experimental efforts, the largest proportion of patients worldwide lives in the chronic phase, where neurological deficits, neuropathic pain, and secondary complications persist for decades. According to the World Health Organization (WHO) (38), an estimated 15.4 million people were living with an SCI in 2021, with a substantial burden of long-term disability and reduced life expectancy. Epidemiological analyses confirm that SCI disproportionately affects young adults, primarily men between 30 and 40 years of age, resulting in a long survival window during which chronic symptoms accumulate and quality of life is profoundly impacted (1,2,3).

Individuals living with chronic SCI frequently develop persistent NeP, affecting up to 80% of patients, alongside musculoskeletal pain, spasticity, osteoporosis, pressure injuries, autonomic dysfunction, and metabolic fragility. These health challenges are compounded by social and economic barriers, including reduced employment opportunities, limited access to rehabilitation, and healthcare inequities that contribute to shortened life expectancy in many regions (38).

Despite these obstacles, patients often maintain meaningful personal, social, and professional lives, underscoring the urgent need for interventions capable of acting in the chronic phase, long after the initial trauma. In this context, treatments targeting glial dysfunction, neuroinflammation, synaptic imbalance, and impaired plasticity offer a promising translational avenue. Our efforts would address this clinical gap by evaluating whether BoNT/A, delivered during the chronic post-injury window and combined with EMS, can modulate injury-associated glial responses and restore cellular homeostasis within the chronically injured spinal cord.

This study provides preclinical evidence that the combination of early EMS and delayed BoNT/A treatment leads to striking functional recovery from paraplegia in a severe model of chronic SCI.

In the SCI rehabilitation field, activity-based interventions such as locomotor training (including treadmill-based paradigms) aim to engage residual spinal circuitry and promote stepping-like patterns through repetitive sensorimotor input (18). Neuromodulatory strategies, such as epidural electrical stimulation, can restore voluntary locomotor control by reactivating spared spinal networks (39). These approaches primarily target central locomotor circuits and depend on the preservation of functional residual connectivity.

In our severe contusion model, however, this approach did not translate into measurable recovery, as also indicated by our preliminary treadmill training experiments reported in the Supplementary Methods/Figure S2, where BMS scores remained unchanged over the observation period. In this context, we deliberately prioritized EMS as a peripheral, muscle-directed strategy to address an early and major critical point of severe SCI, rapid disuse-driven atrophy and progressive loss of neuromuscular integrity, rather than attempting to directly “train” locomotion in paraplegic animals. By preserving muscle trophism and neuromuscular connectivity, EMS is expected to stabilize the effector system that ultimately executes movement, thereby increasing the likelihood that any centrally acting intervention can be functionally expressed.

Animals receiving this combined approach (EMS + BoNT/A) showed significant restoration of hindlimb motor function, which was not observed in any other group, including those treated with EMS or BoNT/A alone. These findings underscore the importance of an integrative therapeutic strategy capable of both preserving tissue integrity and modulating the chronic neuroinflammatory environment.

Importantly, we demonstrate that a short-term EMS protocol, initiated early after injury, is sufficient to exert long-term protective effects against muscle atrophy, a key limiting factor in chronic SCI recovery. Although EMS does not induce persistent spinal or regenerative effects, its peripheral impact on the stimulated muscles is long-lasting, as evidenced by the sustained mitigation of atrophy and preservation of muscle structure observed even 60 days after stimulation cessation.

Even when discontinued prior to BoNT/A administration, EMS alone preserved muscle structure and function, likely contributing to the success of subsequent interventions. This supports clinical observations that early EMS enhances tissue trophism, perfusion, and microenvironmental quality (40, 41) but goes further by showing that EMS creates a pro-regenerative niche that remains responsive to neuroactive therapies weeks later.

Beyond conventional rehabilitation (6), regenerative strategies aimed at modifying the chronic inhibitory milieu of SCI are increasingly explored. These include enzymatic degradation of inhibitory extracellular matrix components using chondroitinase ABC (42), as well as cell-based therapies designed to replace neuronal and glial populations or provide trophic support (43). Such approaches attempt to overcome the structural and molecular barriers imposed by the chronic glial scar. Within this broader landscape, our findings support a complementary perspective: rather than viewing rehabilitation and regeneration as parallel or competing strategies, an early conditioning phase that preserves peripheral targets may enhance the effectiveness of later microenvironment-modifying interventions. By mitigating excitotoxicity, inflammation, and astroglial hypertrophy, BoNT/A may contribute to a more permissive lesion environment that could, in principle, synergize with pro-regenerative approaches.

Crucially, BoNT/A was ineffective when administered alone during the chronic phase, failing to restore motor function or reduce tissue degeneration in the absence of EMS preconditioning, unlike when injected during the acute phase (12, 13). This suggests that BoNT/A’s effectiveness in the chronic phase relies on the preservation of structural connectivity and cellular responsiveness, which are supported by EMS. Once these conditions are met, BoNT/A appears to act as a potent disease-modifying agent, reducing gliosis, excitotoxicity, apoptosis, and promoting oligodendrocyte survival and remyelination.

Moreover, we demonstrated that BoNT/A treatment also when administered in chronic phase, in mild/moderate contused mice, can mitigate pain-related mechanisms, significantly attenuating neuropathic pain-like behaviours. While the peripheral analgesic properties of BoNT/A are well-established (18, 43, 44) our findings extend its relevance to central injury models, indicating a broader therapeutic spectrum that includes central antinociceptive activity, likely mediated by glial modulation and rebalancing of excitatory neurotransmission

To elucidate the mechanisms by which BoNT/A promotes motor recovery and alleviates NeP, we demonstrate that BoNT/A induces a profound remodelling of astrocytes in the injured spinal cord, an effect that goes well beyond simple attenuation of astrogliosis. Astrocytes play a pivotal role in regulating glutamate clearance, modulating both excitatory and inhibitory neurotransmission, and maintaining overall synaptic homeostasis (46). Our findings indicate that BoNT/A modulates injury-associated astrocytic reactivity in the chronically injured spinal cord, neurotoxic profile toward a morphology and molecular signature more compatible with homeostatic and neuroprotective states (47, 48, 49). Rather than producing a uniform suppression of gliosis, BoNT/A acts preferentially within the lesion epicenter, the region of highest metabolic burden and glutamate overload, where astrocytes typically display the most pronounced A1-like features (33, 34). This region-specific effect suggests that BoNT/A targets the cellular compartments under maximal stress, possibly by modulating cytoskeletal dynamics and intracellular trafficking, two processes tightly linked to astrocytic hypertrophy (33).

The structural remodeling observed in BoNT/A-treated animals, characterized by compact arbors and reduced territorial expansion, is consistent with a shift away from A1-like morphology, which is defined by extensive process hypertrophy and increased GFAP polymerization. This interpretation is reinforced by molecular signatures known to distinguish astrocytic phenotypes. A1 astrocytes downregulate glutamate transporters and exacerbate inflammatory signaling (32, 33), while A2 astrocytes display enhanced EAAT1/GLAST expression, supporting synaptic function and neuronal survival (33). The preservation of EAAT1 we observed after BoNT/A treatment aligns closely with this A2-like profile and is consistent with prior descriptions of EAAT1 as a hallmark of reparative astrocytes (33).

The modulation of glutamatergic markers further supports this interpretation. Although vGLUT1 (31) remains chronically reduced after injury (50), an established long-term consequence of excitatory circuit remodeling, BoNT/A decreased GFAP–vGLUT1 colocalization, suggesting a more selective and compartmentalized astrocyte–synapse interaction. Importantly, this reduction occurs in the context of preserved EAAT1, indicating that BoNT/A does not impair glutamate buffering but may limit maladaptive perisynaptic enwrapping typical of A1-like astrocytes. A less intrusive, more spatially confined astrocytic presence may help restore synaptic homeostasis and reduce chronic excitotoxic stress.

Together, these observations support the view that BoNT/A promotes a region-selective modulation of astrocytic structure and associated molecular markers. While the present study does not directly define astrocyte “states”, the convergent morphology- and marker-based data are consistent with a shift toward less hypertrophic, functionally supportive astrocytic profiles. This change may arise from both direct modulation of astrocytic signaling pathways and indirect effects on the progenitor cell niches that contribute to astroglial turnover, as suggested by our previous demonstration that BoNT/A enhances the expansion of Nestin⁺ populations in the injured spinal cord (12). In the chronic phase of SCI, where gliosis becomes maladaptive and contributes to synaptic dysfunction, inflammatory persistence, and metabolic impairment, the ability of BoNT/A to remodel astrocytic structure and restore key homeostatic functions may represent a mechanism underlying its therapeutic potential.

The observed modulation of microglial cell morphology and phenotype following BoNT/A treatment points to a broader anti-inflammatory mechanism beyond astrocytic remodelling. Chronic activation of microglia has been recognized as a key driver of secondary degeneration in spinal cord injury, sustaining a pro-inflammatory milieu through persistent release of cytokines (e.g., IL-1β, TNF-α), reactive oxygen species, and matrix metalloproteinases (51, 52).

Importantly, this study captures a cellular landscape that remains poorly characterized in the literature, the chronic phase of SCI, both because most animal models fail to fully mirror long-term human pathology (14, 53) and because severe contusion models rarely include extended temporal analyses of glial dynamics (54).

Our data reveal an unexpectedly heterogeneous and spatially segregated glial environment in chronic SCI, where distinct microglial, astrocytic, and oligodendroglial phenotypes (discussed below) coexist in region-specific patterns. This complexity underscores the importance of studying chronic time points, where tissue remodeling, ongoing degeneration, and compensatory responses converge into a highly dynamic neuroimmune microenvironment.

In the chronic phase of SCI, microglia display substantial morphological heterogeneity, which likely reflects the intrinsic complexity of long-term neuroinflammation rather than experimental variability. This diversity of phenotypes, coexisting within the same tissue and differing across regions, may provide a more faithful representation of the heterogeneous cellular responses reported in clinical SCI. Within this framework, the trends observed in our study suggest that BoNT/A may contribute to maintaining a greater proportion of ramified, less reactive microglia, particularly in the perilesioned area, whereas saline-treated animals more frequently exhibit reactive and rod-like forms typically associated with persistent inflammatory activation.

The general pattern of morphometric analyses such as Sholl profiling is consistent with these phenotypic trends and with the expected variability of chronic SCI, where regional microenvironments strongly shape microglial structure. Similarly, changes in Iba1+ area likely reflect differences in phenotype composition rather than absolute changes in activation, ramified cells occupy larger territories, while reactive forms are more compact despite being more inflammatory.

Overall, although subtle, these converging observations indicate that BoNT/A may modulate microglial behaviour within a highly heterogeneous chronic landscape, supporting a shift toward states less associated with sustained inflammation. Crucially, the variability observed across all groups highlights a fundamental characteristic of chronic SCI and underscores the need to interpret glial responses within this inherently diverse biological context.

These effects echo BoNT/A’s known actions in peripheral inflammatory models, where it suppresses the release of pro-inflammatory mediators from both microglia and macrophages (11, 55). By reducing microglial reactivity at structural and functional levels, BoNT/A may dampen the chronic inflammatory loop that impedes regeneration in SCI.

These morphological changes were paralleled by a selective increase in β-actin expression in the EMS+BoNT/A group, as revealed by Western blot analysis. Although traditionally used as a housekeeping control, actin levels are known to fluctuate in response to cytoskeletal remodelling and glial activation states (46). The observed increase may reflect an ongoing structural reorganization of glial cytoskeleton, supporting the notion that BoNT/A exerts its effects by reprogramming glial morphology and function at both cellular and molecular levels.

Taken together with the astrocytic remodelling described above, the data suggest that BoNT/A orchestrates a coordinated reprogramming of the glial landscape, acting on both astrocytes and microglia, to mitigate chronic neuroinflammation. This dual action likely contributes to the creation of a more permissive environment for neuronal survival and synaptic preservation, reinforcing the therapeutic relevance of BoNT/A as a glia-targeting agent in spinal cord injury and other neuroinflammatory conditions.

Together, these findings suggest that BoNT/A can reshape the chronic SCI environment to promote recovery by acting primarily on glial components. Unlike classical neurotrophic or anti-inflammatory drugs, BoNT/A does not target neuronal receptors or broadly suppress inflammation. Instead, it enables selective glial reprogramming and preservation of oligodendrocyte-mediated myelination, facilitating long-term improvements in tissue integrity and function.

A key outcome of BoNT/A treatment in our chronic SCI model was the preservation of neural tissue integrity, characterized by a significant reduction in apoptotic cell death and an increase in neuronal and oligodendroglia survival. TUNEL and Caspase-3 staining revealed fewer apoptotic cells in BoNT/A-treated animals, with clear evidence of protection in NeuN⁺ neurons and Olig1⁺ oligodendroglia cells. Interestingly, these effects occurred in the absence of synaptic receptor normalization: both NMDA and GABA-A receptor subunit expression remained equally dysregulated across all injured groups, including BoNT/A-treated animals. This further supports the idea that BoNT/A operates upstream of direct synaptic repair mechanisms, likely through the modulation of glial-derived signals and perisynaptic homeostasis.

Interestingly, the neuroprotective effects of BoNT/A occurred despite persistent downregulation of both NMDA and GABA-A receptor subunits at 60 dpi. For GABA-A, this aligns with evidence that inhibitory failure after SCI is driven less by receptor abundance and more by interneuron loss, reduced GAD65/67, increased uptake, and chloride dysregulation (KCC2 downregulation/NKCC1 upregulation). BoNT/A likely preserves neurons and oligodendrocytes by reducing excitotoxic and glial-derived stress rather than directly reinstating GABAergic tone, suggesting potential synergy with strategies that restore inhibition (e.g., baclofen, GAT inhibitors, KCC2 enhancers) (54–57). For NMDA receptors, chronic dysregulation reflects complex, subunit- and compartment-specific remodelling not captured by segmental Western blots. BoNT/A dampens excitotoxicity upstream but does not normalize NMDA subunit expression, indicating that combinations with NMDA-targeted interventions (e.g., NR2B antagonists, low-dose NMDA blockers, PSD-95/nNOS disruptors) may further enhance functional recovery (60–63).

These findings suggest that BoNT/A engages a multi-cellular neuroprotective program that extends throughout glial modulation to promote neuronal survival.

This neuroprotection may be partly explained by BoNT/A’s capacity to reduce extracellular glutamate (64–66) and attenuate excitotoxic signalling, which are key contributors to secondary degeneration in SCI. By dampening glial reactivity and restoring perisynaptic organization, BoNT/A likely promotes a permissive environment for synaptic reorganization, axonal sprouting, and endogenous repair processes. These effects may help initiate functional improvements such as motor recovery, which has been shown to correlate with reduced neuronal apoptosis and glutamate spillover in chronic SCI (67, 68).

One of the most striking outcomes was the enhanced preservation of oligodendroglia lineage cells and increased myelin protein expression in BoNT/A-treated animals. In ex-vivo, we observed a higher density of Olig1⁺ cells and significant upregulation of MBP. An additional, treatment-independent observation emerging from our OLIG1 analyses concerns the marked phenotypic heterogeneity of oligodendrocyte lineage cells across different anatomical regions of the chronically injured spinal cord. OLIG1-expressing cells adopt distinct morphologies in the epicenter, perilesional areas, and scar tissue, a finding that aligns with developmental and injury-related dynamics previously described in the literature (69). During normal CNS development, OLIG1 undergoes a well-characterized shift from a nuclear to a cytoplasmic localization, a transition required for oligodendrocyte maturation and myelin membrane expansion (70). OPCs also display high morphological plasticity, transitioning from elongated bipolar shapes during early migration to increasingly branched, ramified morphologies as they mature and interact with axons (69,71). In adulthood, OPCs retain a surveying phenotype with stable but dynamic processes, yet upon CNS injury they may either reacquire a hypertrophic, NG2-upregulated profile, or revert to a bipolar motile phenotype to facilitate recruitment toward the lesion core (69).

In vitro, BoNT/A promoted the differentiation of OPCs into mature oligodendrocytes, increased process complexity, and elevated expression of myelin-related markers such as MBP, GalC, PLP, and MOG. These effects were observed under both proliferative and differentiating conditions, indicating that BoNT/A acts even in the absence of extrinsic pro-differentiation cues.

One plausible explanation for this sensitivity lies in the differential expression and role of SNARE proteins along the oligodendrocyte lineage. While SNAP25 is typically regarded as a neuron-specific SNARE, its expression has now been confirmed in human oligodendrocytes (72). Our findings of cl-SNAP25 signal in PLP⁺ and MOG⁺ cells, particularly in BoNT/A-treated non-differentiated cultures, suggest that SNAP25 is not only present but also accessible to BoNT/A enzymatic activity in the oligodendroglia lineage.

Moreover, this effect was most prominent in NG2⁺ OPCs under non-differentiating conditions. The highest levels of cl-SNAP25 in non-differentiated cells suggest that BoNT/A preferentially acts on immature cells, possibly because of higher endocytic activity, differential SNARE expression, or increased expression of BoNT/A receptor (SV2). These hypotheses are supported by studies indicating that OPCs, particularly NG2⁺ cells, display a high degree of synaptic interaction with glutamatergic neurons and possess functional postsynaptic specializations, including vesicle release and uptake machinery (73–75).

Notably, NG2⁺ cells are not merely precursors but are now recognized as active regulators of CNS plasticity. Their ability to receive synaptic input, respond to neurotransmission, and regulate axon-glia interaction positions them as dynamic participants in injury and repair (76, 77). The fact that BoNT/A has a more pronounced effect on these cells in a non-differentiated state raises the possibility that early-stage OPCs are “primed” for BoNT/A-mediated modulation, perhaps via SNARE-dependent control of metabolic vesicle delivery or local calcium buffering, as suggested for early myelin assembly (77).

These findings provide a compelling rationale to further explore how BoNT/A interfaces with the unique synaptic-like physiology of NG2⁺ OPCs, which, unlike mature oligodendrocytes, are capable of bidirectional communication with neurons and other glia (78,79). In this framework, BoNT/A may operate not only as a blocker of excitatory stress but also as a modulator of glial vesicle signalling, tipping the balance toward differentiation and myelination.

Moreover, we observed a strongest response to BoNT/A in terms of branching complexity in non-differentiation condition, measured in MBP and MOG stained, and process length, in differentiated condition.

Of note, our findings seem in contrast with those of Chacon-De-La-Rocha et al. (80), who reported a decrease in NG2⁺ OPC complexity after intrahippocampal BoNT/A administration. However, their use of a much higher BoNT/A concentration (1 nM vs 10 pM) and their targeting of a non-lesioned, synaptically dense brain region likely contribute to this discrepancy. Nonetheless, their study supports the notion that BoNT/A can act directly on OPCs in vivo, further validating our mechanistic hypothesis.

Taken together, these results suggest that BoNT/A action on oligodendrocyte lineage cells may operate through a dual mechanism. On one side, our in vitro experiments with purified OPCs provide evidence for a direct effect, supported by the presence of cl-SNAP25 in PLP⁺/MOG⁺ cells and by the enhanced differentiation observed under both proliferative and differentiating conditions. On the other side, in the ex vivo and in vivo context, BoNT/A may also act indirectly by shaping the spinal cord microenvironment. The reduction of excitotoxicity and pro-inflammatory mediators, as shown in our study, likely decreases secondary cell death and creates a permissive milieu that supports survival and maturation of oligodendrocyte lineage cells. Consistently, BoNT/A-treated animals displayed increased oligodendroglial preservation in parallel with reduced apoptosis. We therefore propose that BoNT/A enhances remyelination through a synergistic interplay between direct OPC modulation and indirect neuroprotective effects within the injured tissue

This study also underscores the importance of context and timing in BoNT/A efficacy (all results are summarized in a schematic representation in supplementary figure S10). Only when combined with early rehabilitation (EMS) did BoNT/A achieve meaningful functional outcomes. This has important implications for clinical trial design, indicating that BoNT/A may be most effective in patients undergoing or having completed structured neuromuscular reconditioning. The observed effects on glia and myelin further support the rationale for targeting chronic neuroinflammation and demyelination in SCI and related neurodegenerative disorders.

The current research, along with other scientific evidence published by our team (12, 13, 15), strongly supports future steps toward translating these findings into an upcoming clinical phase of BoNT/A use in SCI patients. This study is part of a broader translational R&D program aimed at repurposing BoNT/A, following a strategic development plan aligned with European regulatory guidelines.

Our approach is also consistent with the objectives of the pilot project *“Repurposing of authorised medicines: pilot to support not-for-profit organisations and academia”*, launched by the European Medicines Agency and the Heads of Medicines Agencies (81). The outcomes of this initiative, expected to be published in 2025, will contribute to the development and implementation of a formal repurposing framework, which will guide our future studies on BoNT/A as a repurposed therapeutic agent.

To enable clinical translation, we are pursuing a targeted development pathway that includes early dialogue with regulatory authorities and the preparation of a comprehensive preclinical dossier to support the initiation of a clinical trial of BoNT/A in SCI patients.

On this basis and by bridging functional rehabilitation and targeted neurobiological modulation, this innovative approach may offer a novel therapeutic avenue where few options currently exist.

### Limitations, Future Directions and Clinical Perspectives

This study demonstrates that BoNT/A can exert robust neuroprotective and pro-regenerative effects in chronic SCI, preserving neurons and oligodendrocytes, reducing apoptosis, and attenuating astrogliosis and microglial activation. At the same time, several aspects deserve further clarification to strengthen both the mechanistic understanding and the translational potential of this approach.

One important aspect concerns the use of female mice. This choice was deliberate, as previously validated in our model (14): male mice often show a degree of spontaneous motor recovery even in the absence of treatment, a feature with little translational relevance given that patients with severe SCI rarely recover motor function spontaneously. Female mice, by contrast, provide a more stable and clinically relevant representation of motor outcome. While this enhances translational fidelity, it also raises the need to investigate sex-specific responses more systematically in future studies, especially as sex differences in neuroinflammation, metabolism, and regeneration are increasingly recognized.

Another point relates to the dosing regimen and follow-up window. We tested a single dose of BoNT/A, chosen in line with its long-lasting biological activity. In humans, therapeutic efficacy typically extends for 2–4 months and can last up to 6 months, consistent with toxin turnover and immune clearance.

As preliminary indications of dosing, the dose of 15 pg/mouse of BoNT/A used in this study can be considered as animal no-observed-adverse-effect-level (NOAEL).

Under this assumption, considering an average weight of mice of 40 gr, our dose corresponds to a mouse NOAEL dose of 0.375x10^-6^ mg/kg and, considering 60 kg as human standard weight, to a human equivalent dose (HED) of 0.033x10^-6^ mg/Kg. Based on the amount of toxin contained in the commercial toxins (80) this HED correspond to 4.5 U Botox/kg, 5 U Dysport/kg and 7.5 U Xeomin/kg, or in other way to 270 U Botox, 300 U Dysport and 450 U Xeomin in a human weighing 60 kg. By reducing these values by a factor of 10 for safety reasons, doses of 27 U Botox, 30 U Dysport, and 45 U Xeomin are obtained, which are doses compatible with the doses currently used in clinic for single administration.

Our 60-day follow-up in mice captures the mid-range of this therapeutic window, but longer time points will be needed to assess whether the beneficial effects persist or decline. Optimizing dosing, timing, and the possibility of repeated administrations will be essential steps toward clinical translation.

Mechanistically, while BoNT/A clearly preserved astrocytic and oligodendroglia populations, the nature of these effects remains to be clarified. The increased astrocytic density associated with reduced hypertrophy suggests a “scar-modulating” rather than “scar-forming” phenotype. Whether this results from proliferation, migration, or phenotypic reprogramming requires targeted analyses, including proliferation markers and lineage tracing as well as changes in the expression of pro-inflammatory mediators, such as cytokines and chemokines, to determine whether the morphological normalization of astrocytes and microglia observed after BoNT/A treatment corresponds to a functional shift toward a less inflammatory state. Similarly, the preservation of oligodendrocyte lineage cells may arise from both direct BoNT/A actions, supported by our in vitro data, and indirect modulation of the microenvironment through reduced excitotoxicity and inflammation. Disentangling these contributions will be a priority for future mechanistic work.

Another limitation is that functional receptor balance was not restored: NMDA and GABA-A receptor subunits remained dysregulated across groups. This finding highlights the complexity of chronic excitatory/inhibitory remodelling after SCI and suggests that BoNT/A acts upstream, reducing excitotoxicity and inflammatory cascades without directly resetting receptor expression or chloride homeostasis. Future studies should therefore include subunit- and region-specific analyses, assessment of KCC2/NKCC1 expression, and functional electrophysiology to evaluate inhibitory and excitatory currents. Importantly, this opens the door to combination strategies, in which BoNT/A could be paired with interventions that directly reinforce GABAergic inhibition or normalize NMDA signalling, maximizing neuroprotection and functional recovery.

Finally, although BoNT/A preserved structural features such as myelination and neuromuscular junctions, functional readouts were not included. Electrophysiological measures of conduction velocity, synaptic efficacy, and network activity will be required to confirm that the structural benefits translate into improved functional connectivity.

Altogether, these results support BoNT/A as a candidate therapeutic in chronic SCI, but they also delineate a roadmap for the next steps: inclusion of both sexes, longer follow-up, optimization of dosing schedules, mechanistic dissection of glial and oligodendroglial responses, and combinatorial approaches that integrate BoNT/A with other pro-regenerative interventions. From a clinical perspective, these directions are particularly relevant, as they mirror ongoing challenges in SCI rehabilitation, sex-specific variability, chronicity of neuroinflammation, limited durability of interventions, and the need for multimodal strategies. By addressing these issues, the translational path for BoNT/A in SCI could move closer to clinical trial readiness

## Conclusions

This study demonstrates that botulinum neurotoxin A, when applied in a context of preserved neuromuscular integrity via electrical stimulation, can exert significant neuroprotective and pro-regenerative effects in a murine model of chronic spinal cord injury. BoNT/A modulates the spinal environment by reshaping astrocytic and microglial reactivity, reducing excitotoxic signalling, preserving oligodendrocyte lineage cells, and promoting remyelination. These actions appear to involve both direct effects on glial cells and indirect modulation of the extracellular milieu.

Our results highlight BoNT/A as a unique therapeutic candidate capable of engaging glial pathways traditionally overlooked in SCI pharmacology. The findings set the stage for mechanistic studies and clinical translation, and advocate for a paradigm shift in the treatment of chronic SCI, from symptomatic management toward glia-focused repair strategies.

## Declarations

### Ethics approval and consent to participate

All procedures were in strict accordance with the European and Italian National law (DLGs n.26 of 04/03/2014, application of the European Communities Council Directive 2010/63/UE) on the use of animals for research (Italian Ministry of Health - protocol code 122/2019PR) and with the guidelines of the Committee for Research and Ethical Issues of IASP.

## Availability of data and materials

The datasets generated and/or analysed during the current study are available from the corresponding author on reasonable request.

## Authors’ contributions

Conceptualization, SM, VM; Methodology, SM, LM, MC, VM, FDS, FDA, CP, FP, SL, OR, GS; Investigation, SM, SL, GR, VR, LAP, CP, VM, VV, FDS, LM, SA, ADE, RM; Statistics, SM, VM, LM, MC; Writing – Original Draft, SM, VM; Writing – Review and Editing, SM, SL, OR, LM, GS; Funding Acquisition, SM, LM

## Acknowledgements

The authors thank Prof. Cesare Montecucco (University of Padua) for kindly providing purified botulinum neurotoxin A. The authors also acknowledge the technical support of the EMMA Infrafrontier/Mouse Clinic facilities at CNR-IBBC Monterotondo.

