## Supplemental Materials for "A Translational Preclinical Strategy for Chronic Spinal Cord Injury: Neuroprotective and Regenerative Potential of Botulinum Neurotoxin Type A combined with Muscle Atrophy Prevention via Electrostimulation"

**Authors’ Affiliations:**

^1^ National Research Council of Italy, Institute of Biochemistry and Cell Biology, 00015 Monterotondo (RM), Italy.

^2^ Department of Anatomical, Histological, Forensic Sciences and Orthopedics, Sapienza University of Rome, 00161 Rome, Italy.

^3^ Laboratory affiliated to Istituto Pasteur Italia-Fondazione Cenci Bolognetti, 00161 Rome, Italy.

^4^ Department of Biological, Geological, and Environmental Sciences, Alma Mater Studiorum University of Bologna, Bologna, Italy

^5^ Cellular Neurobiology Unit, Santa Lucia Foundation, 00179 Rome, Italy

^6^Department of Biomedical Sciences, University of Padova and National Research Council of Italy, Institute of Neuroscience, Padova, Italy

^7^ National Research Council of Italy, Department of Biomedical Science, 00185 Rome, Italy

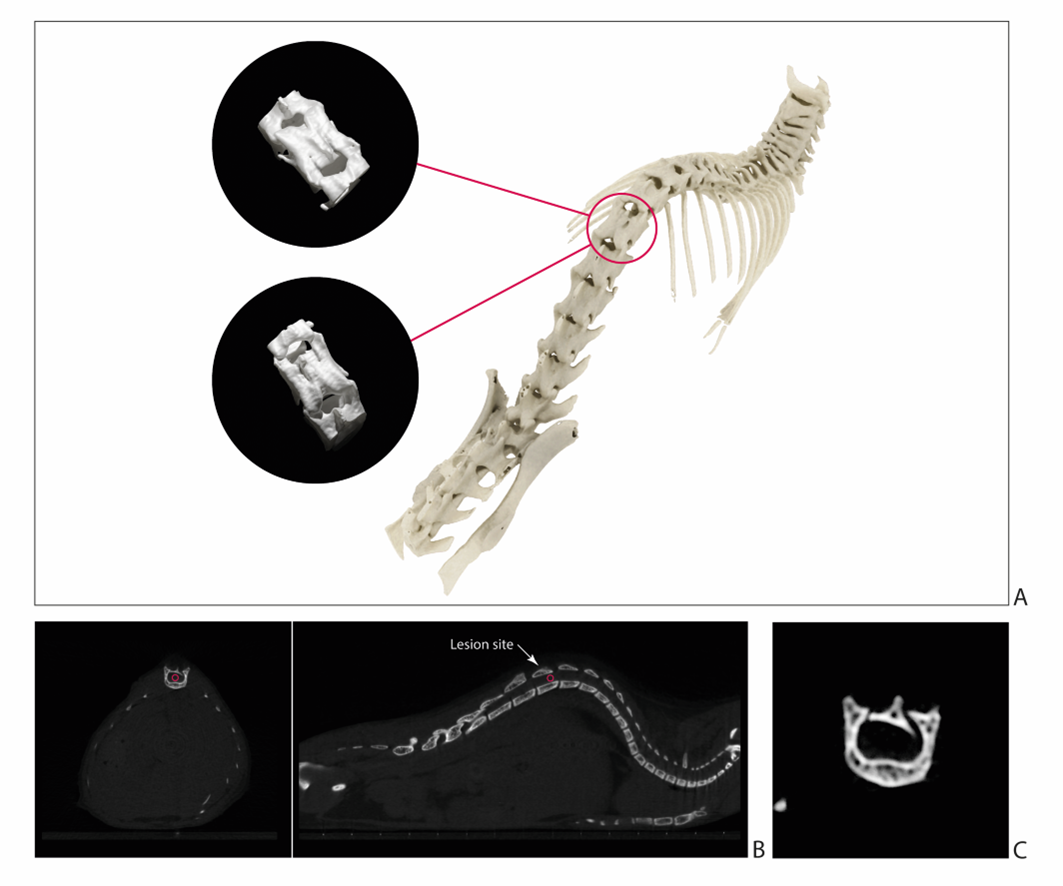

**Figure S1. Micro-CT representative images of a contused spinal cord.** **A)** 3D reconstructed rendering of the thoracolumbar spine, showing the vertebral column and the site of injury. **B)** Axial and Sagittal views of the spinal region highlighting the area of contusion. **C)** A particular of the axial section demonstrating the structural disruption at the lesion site.

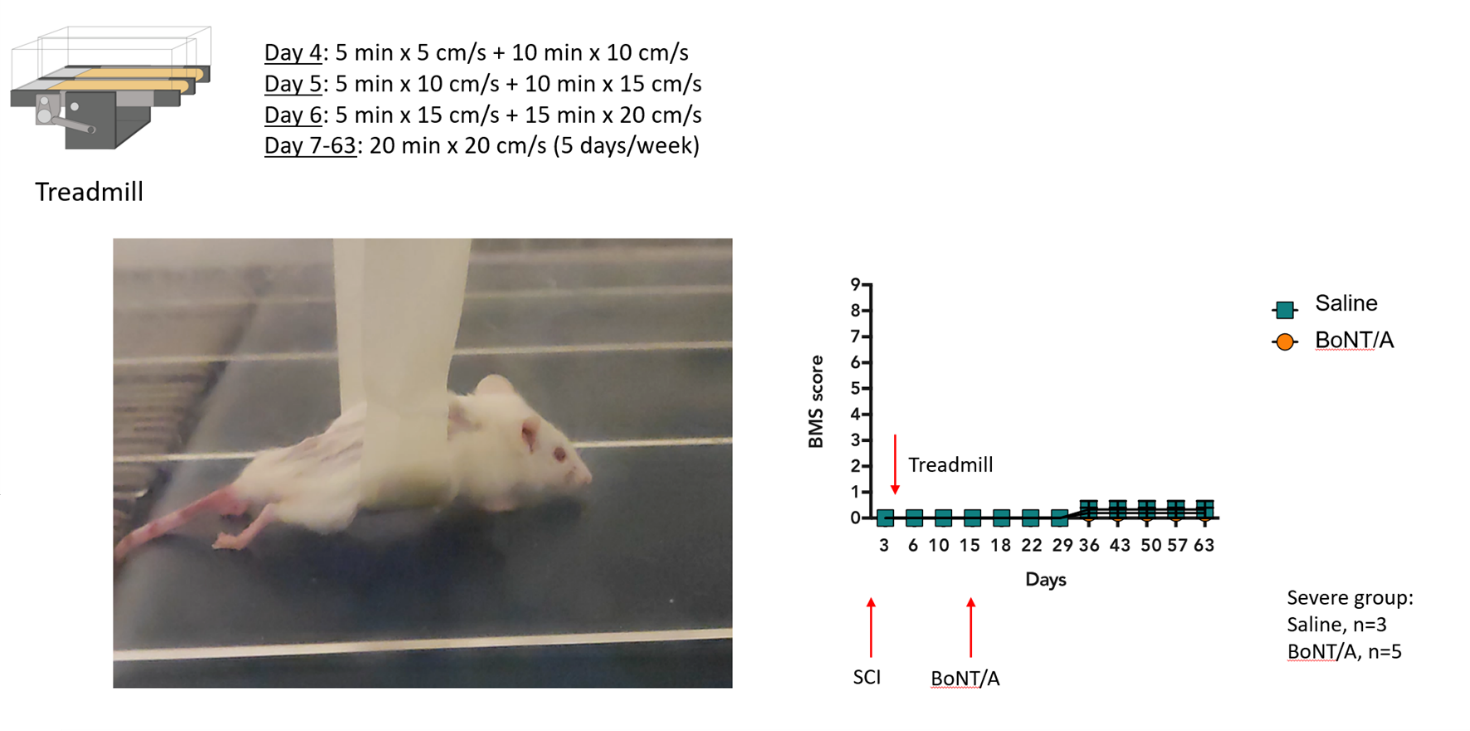

**Figure S2.** **Treadmill training protocol**. To evaluate whether passive locomotor rehabilitation could facilitate recovery in the chronic phase, we tested a treadmill training protocol in severely injured mice. The rationale was to exploit residual spinal locomotor circuits (central pattern generators) that can be activated by repetitive stepping-like movements.

Mice were placed on a motorized treadmill (model LE8710, PanLab, Cornella, Spain) starting 4 days post-injury, and the protocol was progressively intensified over the first week (Cobianchi S, et al. Neuroscience. 2010 doi: 10.1016/j.neuroscience.2010.03.035) followed by maintenance training up to 63 days post-injury:

- **Day 4**: 5 min at 5 cm/s + 10 min at 10 cm/s
- **Day 5**: 5 min at 10 cm/s + 10 min at 15 cm/s
- **Day 6**: 5 min at 15 cm/s + 15 min at 20 cm/s
- **Day 7–63**: 20 min at 20 cm/s, 5 days/week

Despite consistent application of the protocol, treadmill training did not improve locomotor outcomes in the severe SCI group. As shown, Basso Mouse Scale (BMS) scores remained stable at 0–1 throughout the 60-day observation period, with no differences between BoNT/A- and saline-treated animals (Saline, n=3; BoNT/A, n=5). Based on this lack of efficacy, treadmill rehabilitation was not pursued further, and we focused on EMS as a non-invasive and translational strategy to prevent muscle atrophy and enhance responsiveness to BoNT/A treatment.

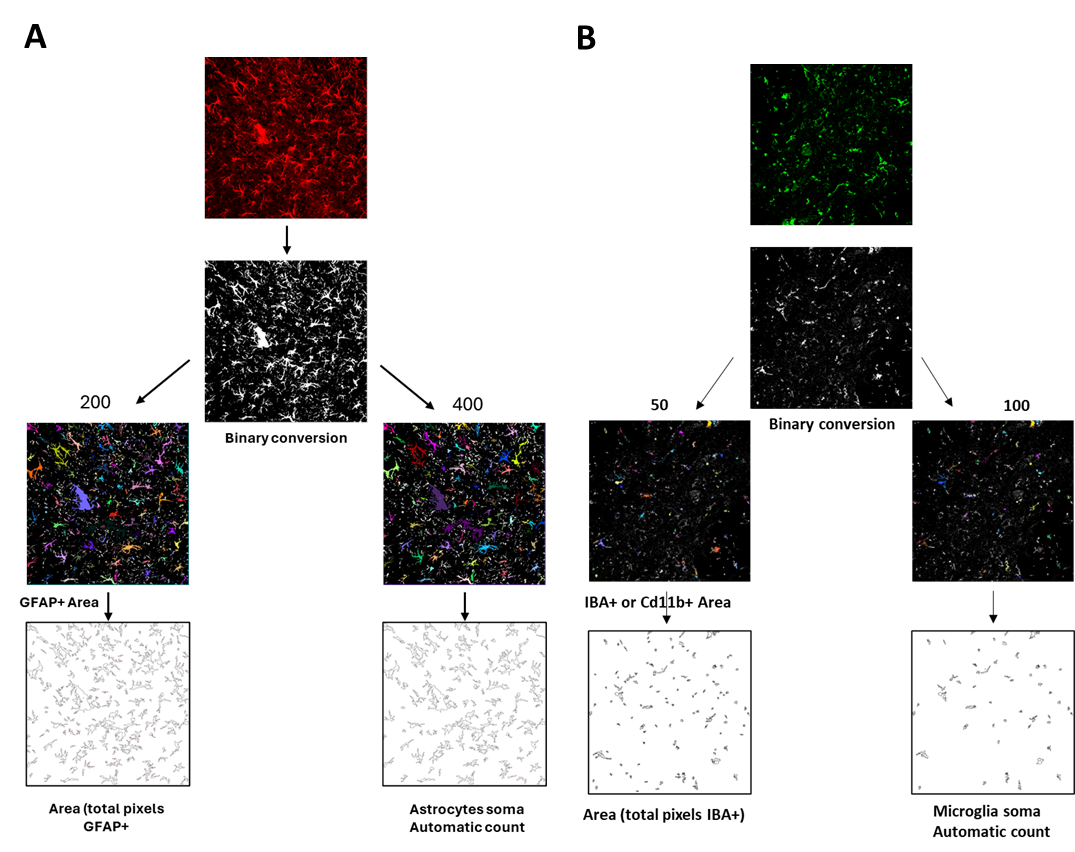

**Supplementary Figure S3. Workflow for astrocyte (A) and microglial (B) quantification analysis.**
Representative sequence of image processing steps used for the automated quantification of GFAP-positive astrocytes (A) and Iba1+/CD11b+ microglia (B). **(A) Astrocytes.** Original confocal image of GFAP immunofluorescence (red) acquired from the spinal cord epicenter (40× magnification, scale bar = 50 µm). The same image converted to 8-bit grayscale and binarized (white = GFAP-positive signal; black = background). Segmentation masks generated using the *Analyze Particles* function in ImageJ, applying two distinct size thresholds (200 px–∞ for total GFAP-positive area; 400 px–∞ for astrocytic somata). Each detected object is pseudo-colored for visualization. Binary outlines showing the individual astrocytic elements detected under each threshold condition. These steps enable automated quantification of both total GFAP-positive area and astrocyte number across spinal cord regions (epicenter, scar, perilesional areas). **(B) Microglia.** Original confocal image of Iba1 or CD11b immunofluorescence (green or red, depending on staining) acquired from the same anatomical regions (40× magnification, scale bar = 50 µm). Image conversion to 8-bit grayscale and binarization to isolate microglia-positive signal. Segmentation of microglial elements using the *Analyze Particles* tool, with size thresholds adjusted to the smaller and more heterogeneous morphology of microglial cell bodies and processes (e.g., 50 px–∞ for total Iba1+/CD11b+ area; 80–100 px–∞ for cell body counts). Each segmented element is automatically assigned a unique color, highlighting the distribution and density of microglial structures. Binary outlines representing individual microglial elements under the two thresholding conditions. This workflow allows automated quantification of both the total microglia-positive area and the number of microglial cells, enabling comparisons across spinal cord regions (epicenter, scar, perilesional), and between experimental groups.

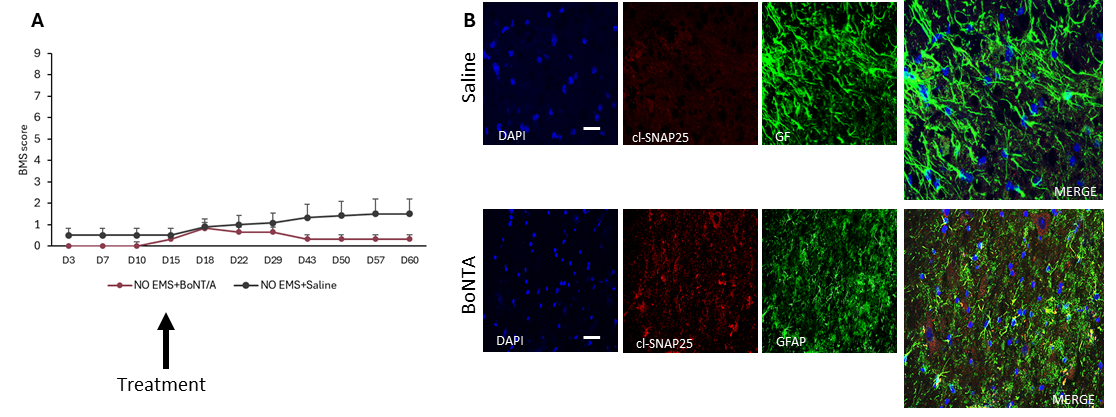

**Figure S4. Spinal administration of BoNT/A or Saline in the chronic phase (15 days post-injury). A)** Motor function was assessed using the Basso Mouse Scale following spinal cord injury. The arrow indicates the time point of intrathecal injection of Saline or BoNT/A. Only animals with severe injury (BMS score between 0 and 3) were included. No significant differences were observed between BoNT/A- and Saline-treated animals (N=6/group; preliminary study). **B)** Representative confocal images (40x magnification; scale bar: 50 μm) of spinal cord tissue near the lesion site (T12–T13), collected 60 days post-injury. Sections were stained for nuclei (DAPI, blue), cleaved SNAP-25 (cl-SNAP25, red), and glial fibrillary acidic protein (GFAP, green). In BoNT/A-treated animals, cl-SNAP25—absent in saline controls—is strongly expressed and partially colocalized with GFAP, indicating sustained toxin activity even 45 days after injection. A reduction in astrocyte activation is also apparent.

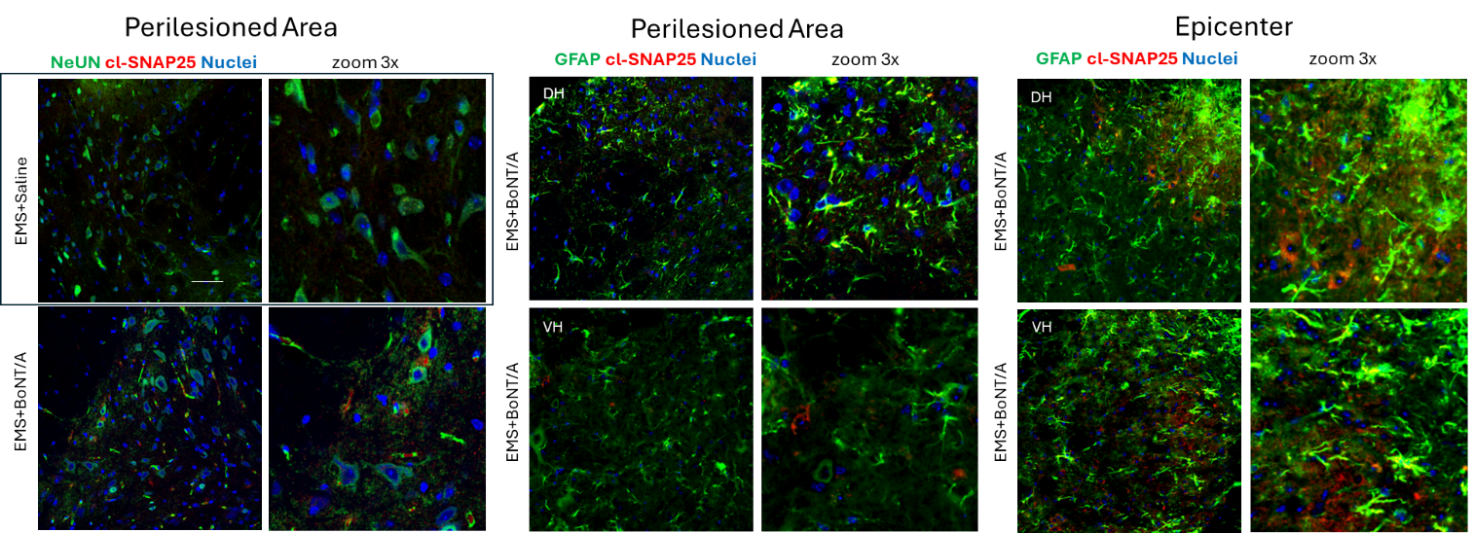

**Figure S5. Detection of cleaved SNAP-25 in spinal cord 60 days post-injury.** Representative immunofluorescence images (40×; scale bar = 50 µm) showing cleaved SNAP-25 (cl-SNAP25, red) in spinal cord sections collected from perilesioned/epicenter areas (T9–T11) 60 days after injury. NeuN (green) was used to label neurons (left panels), and GFAP (green) to label astrocytes (middle and right panels). Nuclei are counterstained with DAPI (blue). The inset on the left shows a saline-treated control section, while all other panels correspond to EMS + BoNT/A-treated animals. The yellow colocalization signal highlights cleaved SNAP-25 within NeuN⁺ or GFAP⁺ cells, confirming the persistence of BoNT/A enzymatic activity in both neuronal and astrocytic compartments. The detection of cleaved SNAP-25 at thoracic levels distant from the lumbar injection site indicates a retrograde transport of the toxin and a long-lasting catalytic action up to 60 days post-administration. DH: dorsal horn; VH: ventral horn.

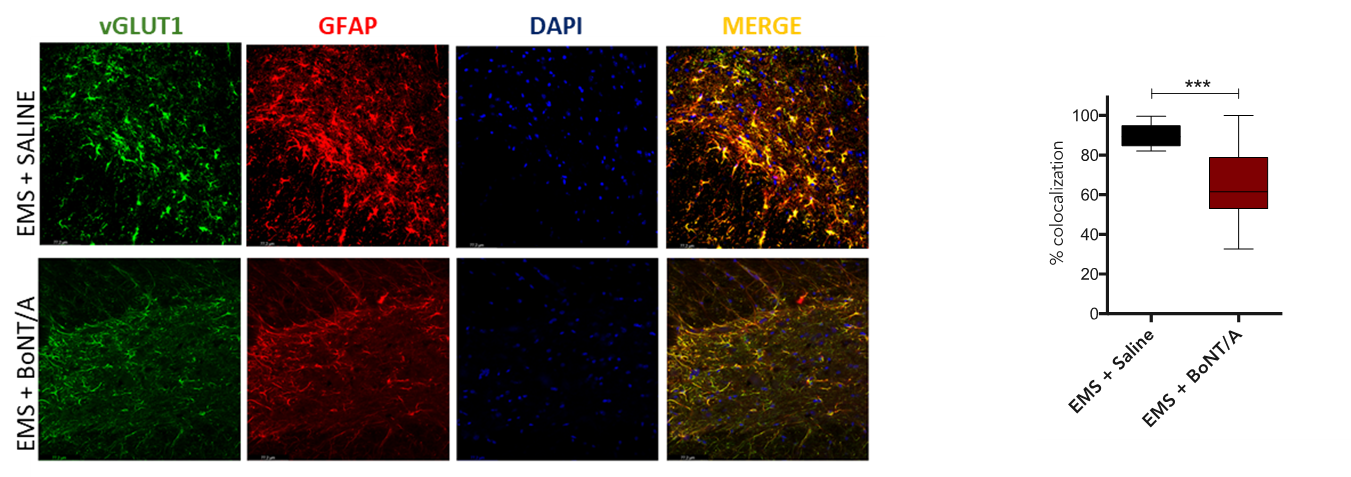

**Figure S6. Immunofluorescence for GFAP and vGLUT1 colocalization analysis.** Representative high-magnification confocal images (40x) from spinal cord sections collected 60 days after SCI, showing individual fluorescence channels for vGLUT1 (green), GFAP (red), and nuclei (DAPI, blue), as well as their merged image. Top row: EMS + Saline group; bottom row: EMS + BoNT/A group. Differential distribution and colocalization of astrocytic processes with the glutamate transporter across treatments. **(D)** Left: Box plot showing percentage of colocalization between GFAP (astrocytes) and vGLUT1 (excitatory presynaptic terminals) 60 days after SCI. BoNT/A significantly reduces GFAP–vGLUT1 colocalization compared to EMS+Saline (Unpaired t-test: t_36_=5.13, ***p<0.0001; values represent mean ± SEM)

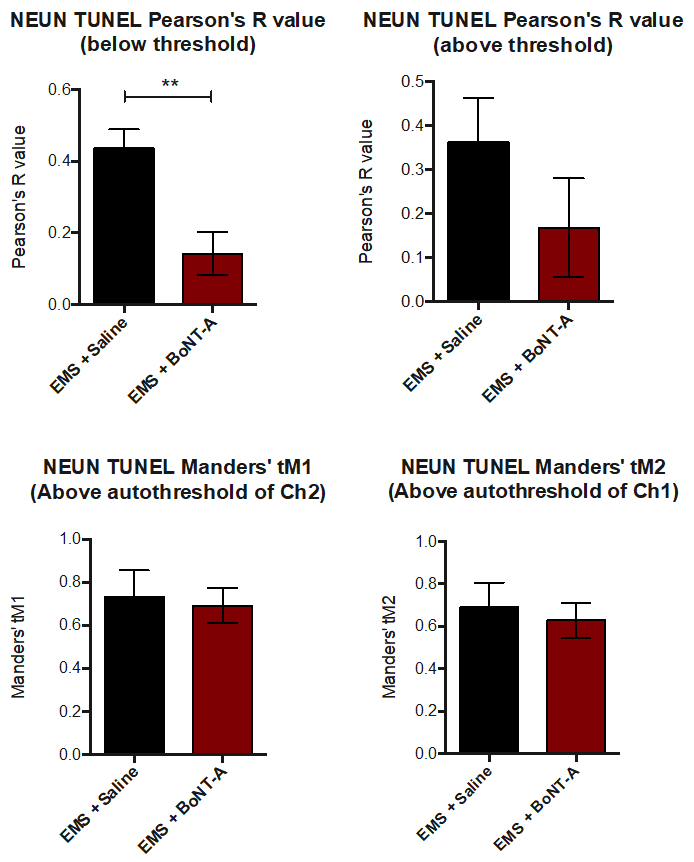

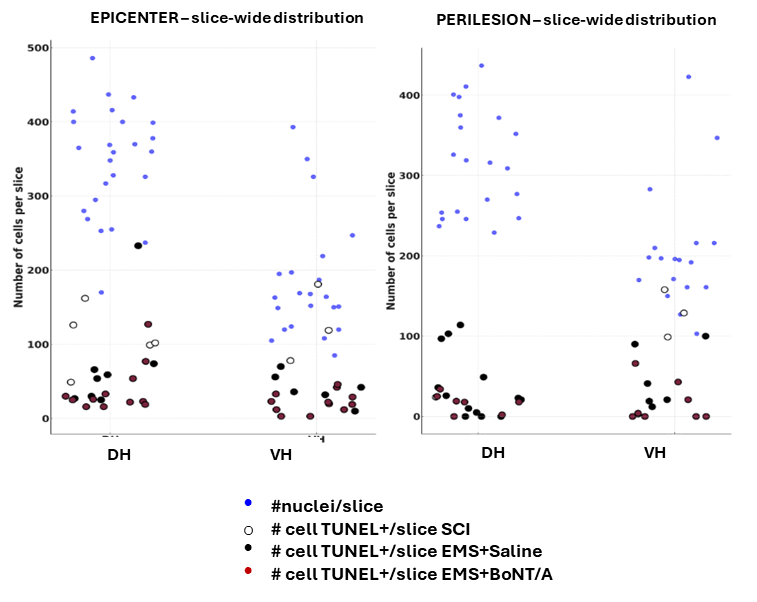

**Supplementary Figure S7.** Quantification of NeuN and TUNEL co-localization in spinal cord sections from EMS+Saline and EMS+BoNT/A groups. **Top panels:** Pearson’s correlation coefficients were calculated below (left) and above (right) the intensity threshold. A significant reduction in the below-threshold Pearson’s R value was observed in the EMS+BoNT/A group, indicating decreased co-localization between NeuN and TUNEL signals (p < 0.01). **Middle panels:** Thresholded Manders’ coefficients (tM1 and tM2), calculated above the automatic threshold for each channel, showed a similar trend but did not reach statistical significance. Data are presented as mean ± SEM (N=3-4 animals/group, 7-14 slices/treatment). **Low panels:** nuclei (blue) and apoptotic cells from different groups distribution across the tissue. Individual values were plotted separately for the dorsal horn (DH) and ventral horn (VH).

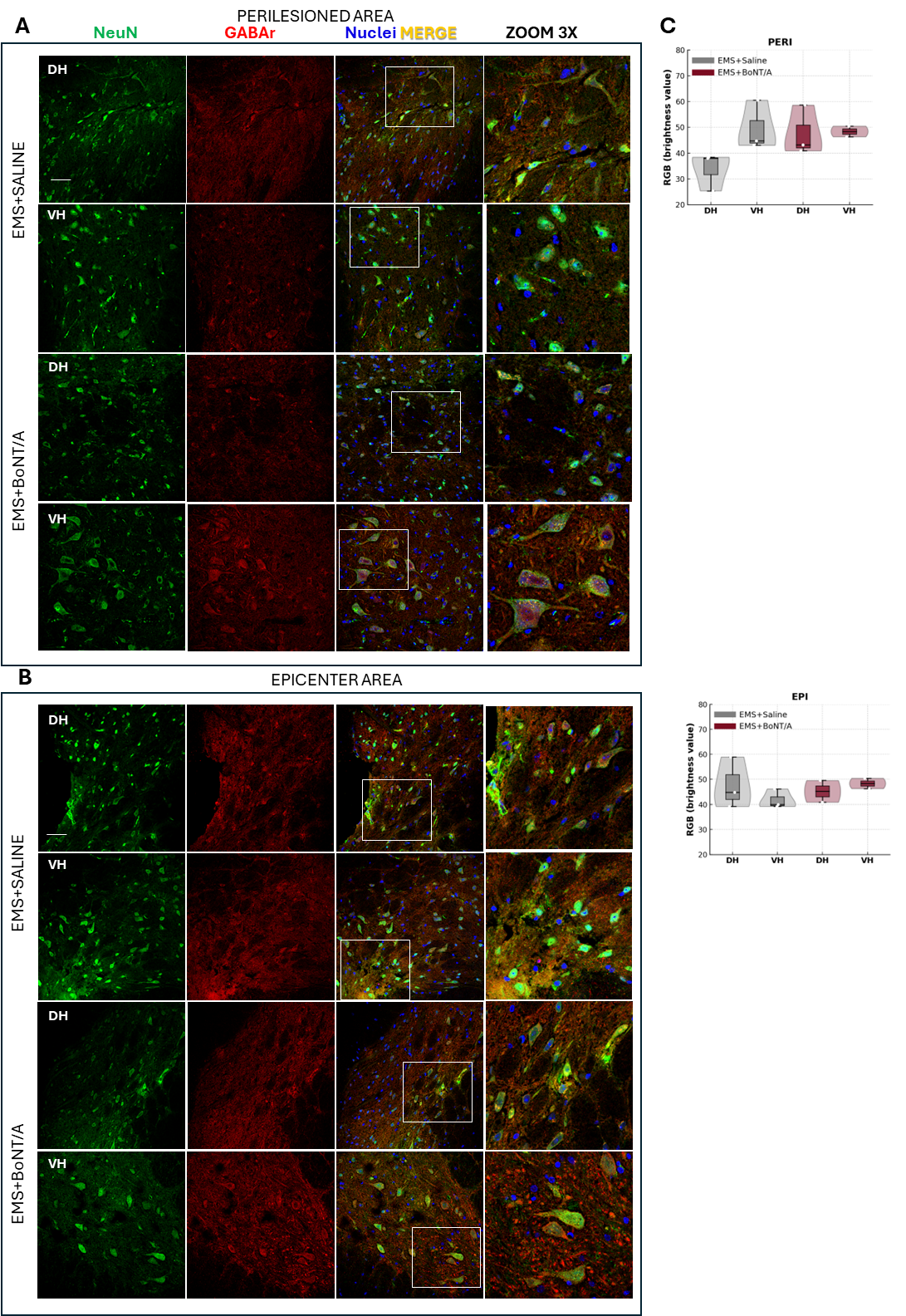

**Supplementary Figure S8. Immunofluorescence analysis of GABA-A receptor α2 in the spinal cord 60 days after SCI.** **(A–B)** Localization and distribution of GABA-A receptor α2 (GABA-Rα2) immunoreactivity in the dorsal (DH) and ventral horns (VH) of the spinal cord from EMS+Saline and EMS+BoNT/A treated animals. **(A)** Representative confocal images of perilesional regions (rostral–caudal, within 2–3 mm from the impact zone; T7–T13 segments). **(B)** Representative images of the epicentral area (T9–T11), corresponding to the region directly affected by the trauma. Sections were co-stained for NeuN (neuronal marker, green), GABA-Rα2 (red), and nuclei (DAPI, blue). Images were acquired at 40× magnification; scale bar = 50 µm. **(C)** Quantitative analysis of fluorescence intensity in DH and VH across epicentral and perilesional regions. Each point represents one animal (*N* = 2–3 per group). All slice-level values (one to two per animal for each area and DH/VH) were averaged per animal, which was considered the experimental unit. Boxplots display median ± IQR, and violin plots illustrate data distribution. Because of the small sample size (*N < 5 per group*), non-parametric descriptive statistics were applied. Kruskal–Wallis and Scheirer–Ray–Hare tests did not reveal significant differences between treatments or areas (*p* > 0.05), indicating that GABA-A receptor expression patterns are comparable between EMS+Saline and EMS+BoNT/A groups

.

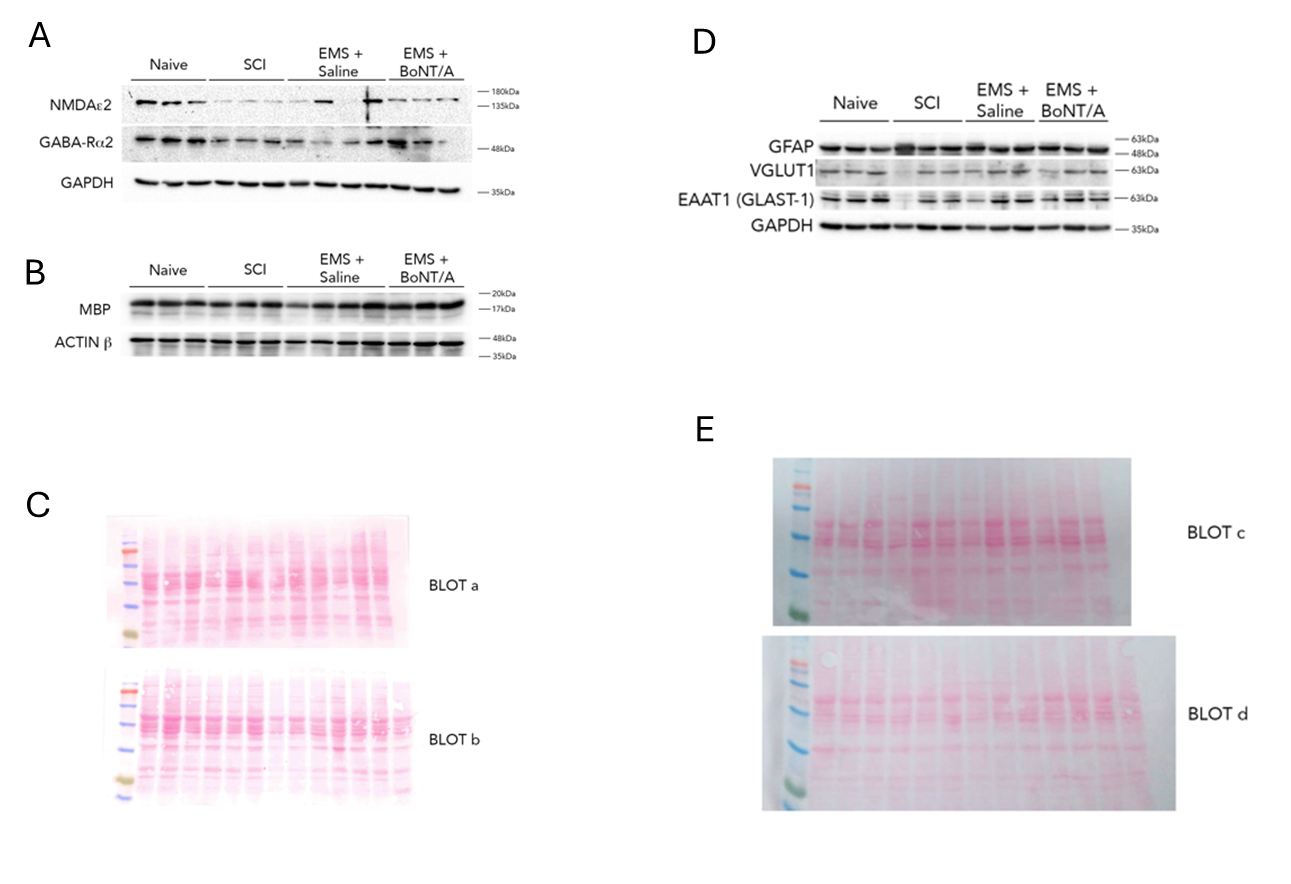

**Figure S9. Full panel of Western blot experiments assessing receptor subunits, myelin integrity, and loading controls in spinal cord tissue 60 days after SCI. A)** Further blots from the same experimental groups (Naïve, SCI, EMS+saline, EMS+BoNT/A) used in Figure 6, showing the expression of: NMDA receptor subunit ε2 (NMDAε2) and GABA-A receptor subunit α2 (GABA-Rα2), associated with excitatory and inhibitory neurotransmission; **B)** Myelin Basic Protein (MBP), a marker of myelin integrity; GAPDH and β-actin, used as reference proteins. Due to variable expression of housekeeping proteins across experimental conditions (see Figure 6 legend), normalization to GAPDH or β-actin was not applied. **C)** Final assessment of protein loading was performed via Ponceau S staining (not shown here, to be included). All samples refer to spinal cord lysates collected 60 days post-injury.

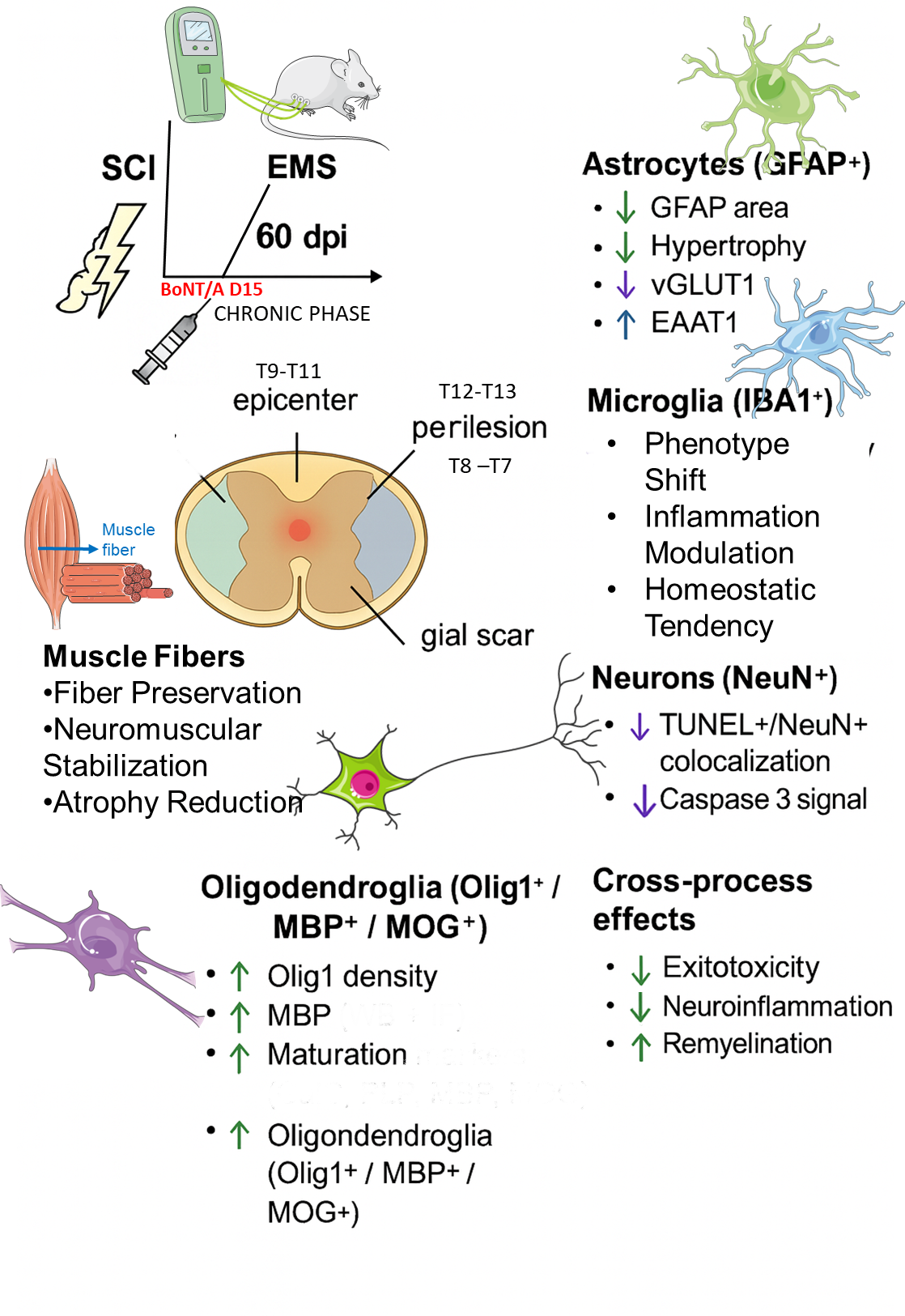

**Figure S10.** Schematic representation illustrates the experimental design and the multi-level effects of combined EMS + BoNT/A treatment administered during the chronic phase of spinal cord injury (SCI). At 60 days post-injury, mice underwent an EMS rehabilitation protocol, with BoNT/A delivered at day 15 of the chronic phase. Histological and molecular analyses were performed in two spinal cord regions, epicenter (T9–T11) and perilesion (T8–T7 / T12–T13), as well as in hindlimb muscles.

The combined treatment modulated several cellular targets:

- Astrocytes (GFAP⁺): reduced hypertrophy and GFAP area, decreased vGLUT1, and increased EAAT1 expression.
- Microglia (Iba1⁺): shift toward less reactive phenotypes, modulation of inflammatory profiles, and increased homeostatic features.
- Muscle fibers: preservation of fiber size, reduced atrophy, and stabilization of neuromuscular junctions.
- Oligodendroglia (Olig1⁺ / MBP⁺ / MOG⁺): increased density, enhanced maturation, and improved myelin-related markers.
- Neurons (NeuN⁺): reduced TUNEL⁺/NeuN⁺ colocalization and decreased caspase-3 activation.

Cross-process analyses indicate reduced excitotoxicity, attenuated neuroinflammation, and enhanced remyelination, supporting the therapeutic potential of EMS + BoNT/A in chronic SCI.

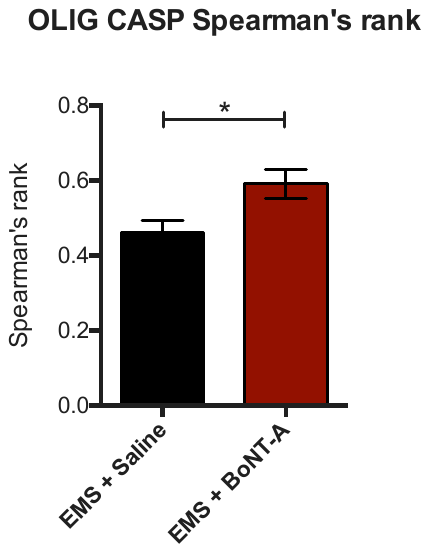

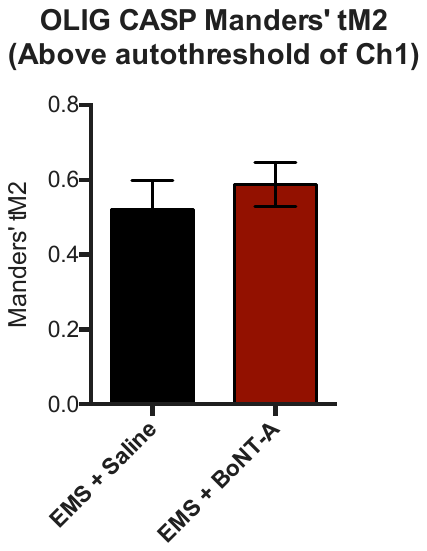
